# Transient mitochondria dysfunction confers fungal cross-resistance between macrophages and fluconazole

**DOI:** 10.1101/2021.01.14.426672

**Authors:** Sofía Siscar-Lewin, Toni Gabaldón, Alexander M. Aldejohann, Oliver Kurzai, Bernhard Hube, Sascha Brunke

## Abstract

Loss or inactivation of antivirulence genes is an adaptive strategy in pathogen evolution. *Candida glabrata* is an important opportunistic pathogen related to baker’s yeast, with the ability to both, quickly increase its intrinsic high level of azole resistance and persist within phagocytes. During *C. glabrata*’s evolution as a pathogen, the mitochondrial DNA polymerase, CgMip1, has been under positive selection. We show that *CgMIP1* deletion not only triggers loss of mitochondrial function and a *petite* phenotype, but increases *C. glabrata*’s azole and ER stress resistance, and importantly, its survival in phagocytes. The same phenotype is induced by fluconazole and by exposure to macrophages, conferring a cross-resistance between antifungals and immune cells, and can be found in clinical isolates despite its slow growth. This suggests that *petite* constitutes a bet-hedging strategy of *C. glabrata*, and potentially a relevant cause of azole resistance. Mitochondrial function may therefore be considered a potential antivirulence factor.

## INTRODUCTION

Human pathogenic fungi remain an underestimated threat in global health, and the mortality rates of fungal infections worldwide are higher or similar to deaths due to malaria or tuberculosis (Bongomin, Gago, Oladele, & Denning, 2017; Kainz, Bauer, Madeo, & Carmona-Gutierrez, 2020). *Candida* species are among the most important human fungal pathogens and cause millions of mucosal and life-threatening systemic infections each year (Bongomin et al., 2017). *Candida glabrata* has become the second most common *Candida* species for immunocompromised patients, surpassed only by *C. albicans* as the primary cause of candidiasis (Lamoth, Lockhart, Berkow, & Calandra, 2018). However, most of the well-characterized pathogenicity mechanisms of *C. albicans* are not shared by *C. glabrata*, and unlike the first, *C. glabrata* does not cause significant host cell damage or elicit strong host immune responses (Brunke & Hube, 2013). Among the main clinical relevant attributes and pathogenic traits of *C. glabrata* are rather a high intrinsic resistance to azole antifungals and an ability to survive for long time and replicate within mononuclear phagocytes (Brunke & Hube, 2013; Gabaldon et al., 2013; Kasper, Seider, & Hube, 2015). Its redundant anti-oxidative stress mechanisms, combined with its ability to modify the phagosomal pH, may partially account for the remarkable ability to survive phagocytosis by macrophages (Cuellar-Cruz et al., 2008; Cuellar-Cruz, Lopez-Romero, Ruiz-Baca, & Zazueta-Sandoval, 2014; Seider et al., 2011). These facts have led to the speculation that *C. glabrata* may take advantage of these immune cells to succeed as a pathogen and disseminate within the host (Kasper et al., 2015).

Among the strategies that confer pathogenicity, the loss or inactivation of certain genes, termed antivirulence genes, is common in pathogenic microorganisms (Bliven & Maurelli, 2012): Cellular pathways and functions that are normally advantageous for the microbe can become superfluous or even disadvantageous under infection conditions, and the loss or inactivation of their encoding genes becomes adaptive during infection. Several examples of such antivirulence factors are known in human pathogenic fungi, and many more are likely to exist (Siscar-Lewin, Hube, & Brunke, 2019).

*C. glabrata* is more closely related to the brewer yeast *Saccharomyces cerevisiae* than to *C. albicans* (Dujon et al., 2004) and clusters with members of the Nakaseomyces group, a genus that includes other environmental and human-associated species (Gabaldon et al., 2013). In a systematic genomic comparison within this group, four genes showed hallmarks of positive selection in *C. glabrata* (Gabaldon et al., 2013). These genes exhibit a relatively high ratio of non-synonymous to synonymous mutations (*d*_*N*_*/d*_*S*_), indicating positive selection during the diversification of *C. glabrata* as a species. Therefore, they might be involved in *C. glabrata*’s specific adaptation to the human host. The gene with the highest *d*_*N*_*/d*_*S*_ ratio (3.40) among them is *CgMIP1*, an orthologue of a mitochondrial DNA (mtDNA) polymerase in *S. cerevisiae* (Gabaldon et al., 2013).

mtDNA encodes subunits of the respiratory complexes, which are involved in the production of ATP during oxidative phosphorylation. The consequences of mtDNA loss have been well described in *S. cerevisiae* and *C. glabrata*, which, unlike other pathogenic yeasts such as *C. albicans* or *Cryptococcus neoformans* (Chen & Clark-Walker, 2000; Toffaletti, Nielsen, Dietrich, Heitman, & Perfect, 2004), are known as *petite*-positive yeasts. The *petite* phenotype due to loss of mitochondrial function is characterized by the namesake small colonies, slow growth, inability to use non-fermentable carbon sources, the activation of the transcription factor *PDR1*, and the upregulation of its targets *CDR1* and *CDR2*, which code for ABC efflux pump transporters (Chen & Clark-Walker, 2000). This upregulation confers high resistance to azole antifungals (Brun et al., 2004; Sanglard, Ischer, & Bille, 2001; Zhang & Moye-Rowley, 2001). Indeed, the *petite* phenotype can be obtained by incubation with high concentrations of azole or ethidium bromide (Bouchara et al., 2000; Goldring, Grossman, Krupnick, Cryer, & Marmur, 1970; Sanglard et al., 2001). Ethidium bromide is known to inhibit mtDNA synthesis and degrade the existing mtDNA (Goldring et al., 1970), but how azoles trigger mitochondria dysfunction is not entirely clear. Azole treatment is known to trigger a temporary loss of mitochondria function (Kaur, Castano, & Cormack, 2004), and the few clinical *petite* strains of *C. glabrata* described so far have been mainly isolated from azole-treated patients (Bouchara et al., 2000; Posteraro et al., 2006). One of these isolates has been further characterized (Ferrari, Sanguinetti, De Bernardis, et al., 2011). Surprisingly, these slow growing isolates showed increased virulence in an animal infection model (Ferrari, Sanguinetti, De Bernardis, et al., 2011). However, when its parental strain was made *petite* by ethidium bromide treatment, its virulence was instead reduced. The same was found in another study using an ethidium bromide-induced *petite* (Brun et al., 2005). Thus, the clinical relevance of the *petite* form is still unclear, and its identification from patient samples may be even hindered by its long generation time.

This study investigates the relevance of the presence and absence of mitochondrial function for *C. glabrata*’s adaptation to the host and its pathogenic potential, as well as potential role *CgMIP1* for switching between *petite* and non-*petite* phenotypes. Deletion of *MIP1* results in *petite* forms, but in contrast to the *S. cerevisiae Scmip1*∆ mutant, *C. glabrata Cgmip1*Δ survives phagocytosis by macrophages significantly better than wild type cells. Importantly, the *C. glabrata petite* phenotype is directly induced in wild type strains by phagocytosis and leads to increased azole resistance, but also *vice versa*, with azole-induced *petites* resisting phagocytosis better. This indicates a clinically important positive feedback between two relevant phenotypes: resistance to macrophages and azoles. The clinical relevance of this phenomenon was further corroborated by the detection of a number of *petite* strains in clinical samples.

## RESULTS

### *MIP1* knock-out mutants of *C. glabrata* and *S. cerevisiae* both show *petite* phenotypes but differ in their survival after phagocytosis

Since the *MIP1* gene of *C. glabrata* seems to have been under selective pressure during the pathogen’s evolution, we investigated its functions in virulence-related scenarios. First, we created a deletion mutant of *CgMIP1* (*Cgmip1*∆) and compared it to a similar deletion of the orthologous gene in *S. cerevisiae* (*Scmip1*∆). *ScMIP1* is known to encode a mitochondrial polymerase (Saccharomyces Genome Database (SGD), www.yeastgenome.org) and therefore, we first checked whether the mutants show the *petite* phenotype. As expected, both *Cgmip1*Δ and *Scmip1*∆ showed phenotype typical for *petite* variants (Brun et al., 2004; Sanglard et al., 2001; Zhang & Moye-Rowley, 2001): small colonies on agar plates and absence of reductive mitochondrial power, absence of mtDNA, and lack of growth in non-fermentable carbon sources (Figure 1). Moreover they showed high expression of the efflux pumps-related genes *PDR1* and *CDR1* (*PDR1* and *PDR5* in *S. cerevisiae*) and high resistance to azoles, again typical for *petite* variants (Figure 1). These results therefore show that, like its *S. cerevisiae* counterpart, *CgMIP1* likely encodes a mitochondrial DNA polymerase, and its deletion triggers loss of mtDNA, loss of mitochondrial function, and a *petite* phenotype in both species.

**Figure 1.**
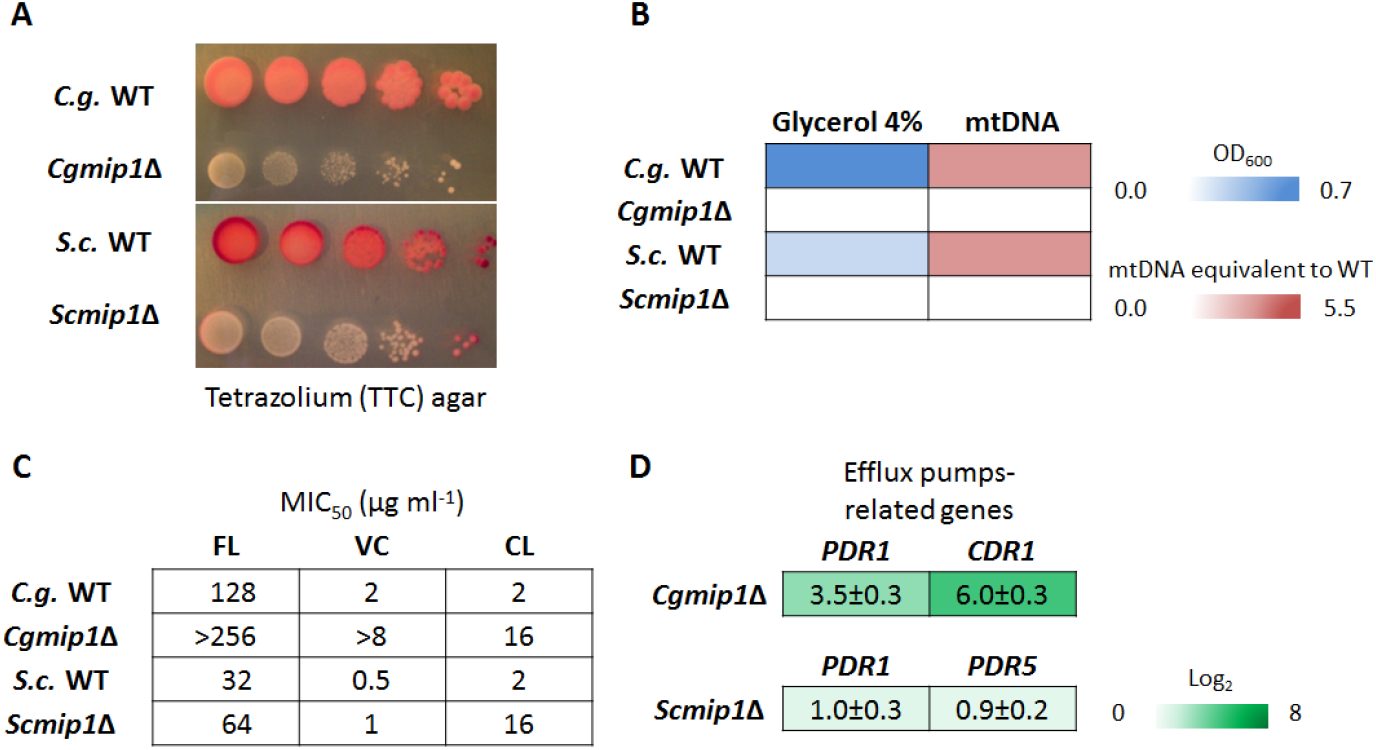
Both *Cgmip1*∆ and *Scmip1*∆ show typical *petite* phenotypes. **(A)** Small colonies and loss of mitochondrial reductive power, as indicated by lack of tetrazolium dye reduction, **(B)** lack of growth in alternative carbon sources like glycerol and absence of mitochondrial (mt) DNA as determined by optical density and qPCR (n=3 for each type of experiment, color by mean), **(C)** high resistance to azoles, including fluconazole (FL), voriconazole (VC), and clotrimazole (CL), and **(D)** overexpression of efflux pumps-related genes (mean ± SD, n=3 independent experiments with 3 technical replicates each).

To study a possible involvement of *MIP1* in processes relevant for virulence, we subjected *Cgmip1*Δ to phagocytosis by human monocyte-derived macrophages (hMDMs) and analyzed its survival after three and six hours. At those time points, macrophages were lysed, and total colony forming units (CFU) were counted after plating on YPD agar. A significantly higher survival rate of *Cgmip1*Δ was found at both times compared to both the wild type control and to *Scmip1*Δ (Figure 2). In order to confirm that this increased number of surviving intracellular yeasts was indeed due to better survival and not due differences in phagocytosis rate or internal replication, we measured both parameters. For phagocytosis rates, *Cgmip1*Δ and wild type were incubated with hMDMs for 1 hour and CFUs from supernatant and macrophage lysate were determined. *Cgmip1*Δ cells were taken up at a slightly higher rate as compared to the wild type (Figure 2), which, however, alone cannot explain the stark increase in the number of intracellular *Cgmip1*Δ especially at six hours: Whereas the *petite* mutant is taken up 1.5 times more than the wild type, the survival of this mutant is up to four times more than the wild type after 3 hours, and six times more at 6 hours.

**Figure 2.**
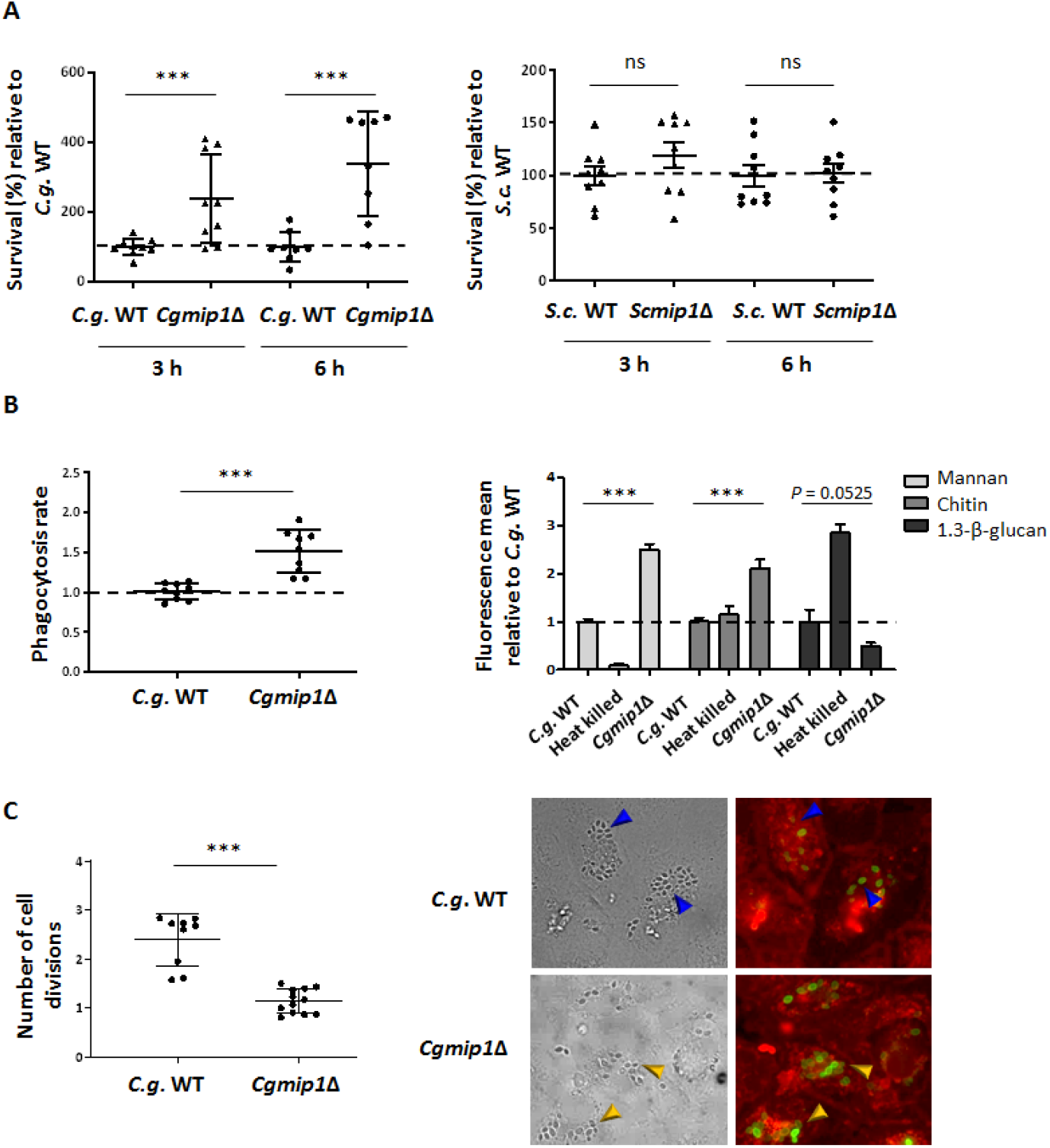
*C. glabrata* and *S. cerevisiae petite* phenotypes differ in their survival after phagocytosis. **(A)** *Cgmip1*∆ survives phagocytosis by hMDMs much better than its parental wild type at early time points up to 6 hours – in contrast to *Scmip1*Δ, which does not show any change in survival compared to its wild type (mean ± SD, n=9 with 3 different donors in 3 independent experiments, each point represents the mean of 3 technical replicates). **(B)** *Cgmip1*∆ is taken up at a higher rate than the wild type by macrophages (mean ± SD, n=9 with 3 different donors in 3 independent experiments, each point represents the mean of 3 technical replicates), and its accessible cell wall structures differ from the wild type (mean ± SD, n=3 independent experiments with 3 technical replicates each). **(C)** In contrast to the wild type, *Cgmip1*∆ does not replicate within the phagosome as shown by FACS (left) and by the lack of FITC-unstained daughter cells (right). These are present in the wild type (blue arrows) contrary to the mutant, which shows only mother cells (yellow arrows) (representative picture shown). Quantitative data is mean ± SD, n=12 with 3 different donors in 4 independent experiments, each point represents the mean of 3 technical replicates. **(A-C)** Statistically significantly different values (unpaired, two-tailed Student’s t-test on log-transformed ratios) are indicated by asterisks as follows: ***, p ≤ 0.001.

To gain more insight into the underlying reason for this slightly increased uptake of *Cgmip1*Δ, the exposure of cell wall components was measured by flow cytometry. Significantly higher exposure of mannan and chitin was observed (Figure 2), while exposure of β(1→3)-Glucan was slightly reduced. Higher surface mannan levels on yeast cells are known to increase phagocytosis rate (Keppler-Ross, Douglas, Konopka, & Dean, 2010), and thus, our results show that mitochondria dysfunction by deletion of *CgMIP1* affects cell wall composition in *C. glabrata* – in agreement with previous observations (Batova et al., 2008; Brun et al., 2005) – and leads to changes in the phagocytosis rate. To also directly measure fungal replication within the macrophages, yeasts were FITC-stained and incubated with hMDMs for six hours. This stain is not transferred to daughter cells, and we measured FITC-negative cells in the macrophage lysate by flow cytometry and also visualized them with fluorescence microscopy. According to our FACS data, *Cgmip1*Δ showed a much lower replication rate than its parental strain, and we did not observe any unstained daughter cells by microscopy. This is in accordance with the nutrient limitation in the phagosome, where only non-fermentable (and therefore *petite*-inaccessible) carbon sources (carboxylic acids, amino acids, peptides, N-acetylglucosamine, and fatty acids) are thought to be available (Gilbert, Wheeler, & May, 2014; Lorenz, Bender, & Fink, 2004; Sprenger, Kasper, Hensel, & Hube, 2018).

These results indicate that, although *Cgmip1*Δ *petite* phenotype is engulfed faster and is largely unable to replicate inside the macrophage, it is killed significantly slower in the first hours of immune cell interaction, in clear contrast to non-pathogenic *S. cerevisiae*.

### The *Petite* phenotype emerges from the wild type after phagocytosis

Our data so far indicates a selective advantage of the *petite CgMIP1* deletion strain during initial interactions with macrophages, despite its inability to replicate within these cells. We therefore analyzed survival of *Cgmip1*Δ and the wild type during long-term residence within macrophages. For this experiment, yeasts were first incubated with macrophages for three hours, the supernatant was removed and the yeast-containing macrophages were then incubated for 7 days. Fungal survival was assessed by CFU counting from plated lysate at 3 hours, 1 day, 4 days, and 7 days. *Cgmip1*Δ again showed higher survival up to one day of co-incubation, in full support of our previous results (Figure 3). However, at later time points, *Cgmip1*Δ showed a significant decrease in survival. This may be explained by its inability to replicate within the phagosome, leading to a long-term disadvantage. Unexpectedly, during incubation of the wild type, we spotted small colonies at all time points, especially after three hours and one day (the time points with the best survival of the *petite* strain), with an average frequency of 1.5×10^−2^ after 1 day (Figure 3). These colonies showed stable small colony formation and lack of growth in glycerol, typical features of *petite* phenotype. Importantly this frequency was higher than the spontaneous emergence of *petite* without macrophages at the same time points (Figure 3). In addition, some of the colonies which did not grow in glycerol gave rise to respiratory-competent phenotype – *i. e*. returned to their original non-*petite* state – when they were plated again on complex media, with an observed frequency of 5.6×10^−2^.

**Figure 3.**
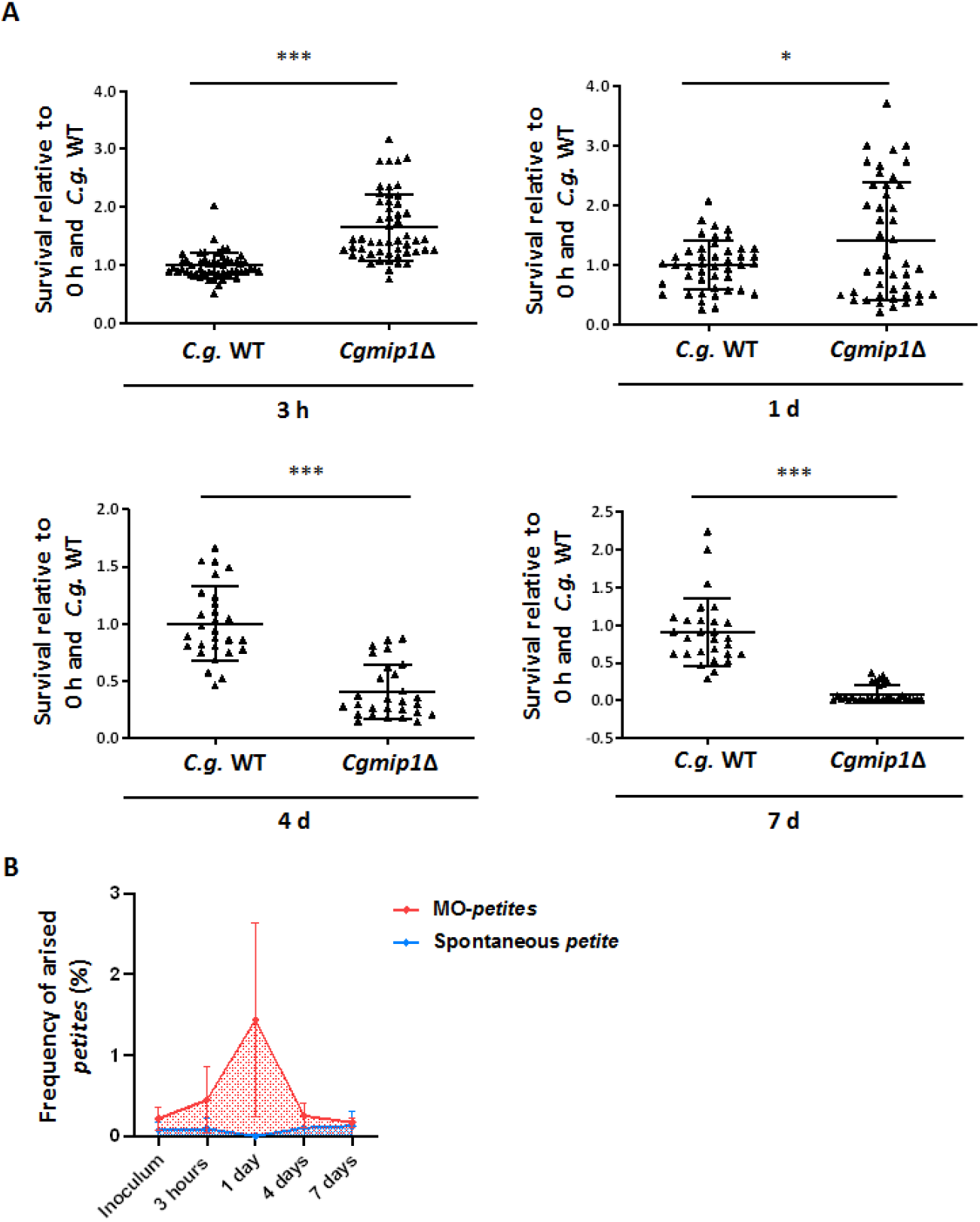
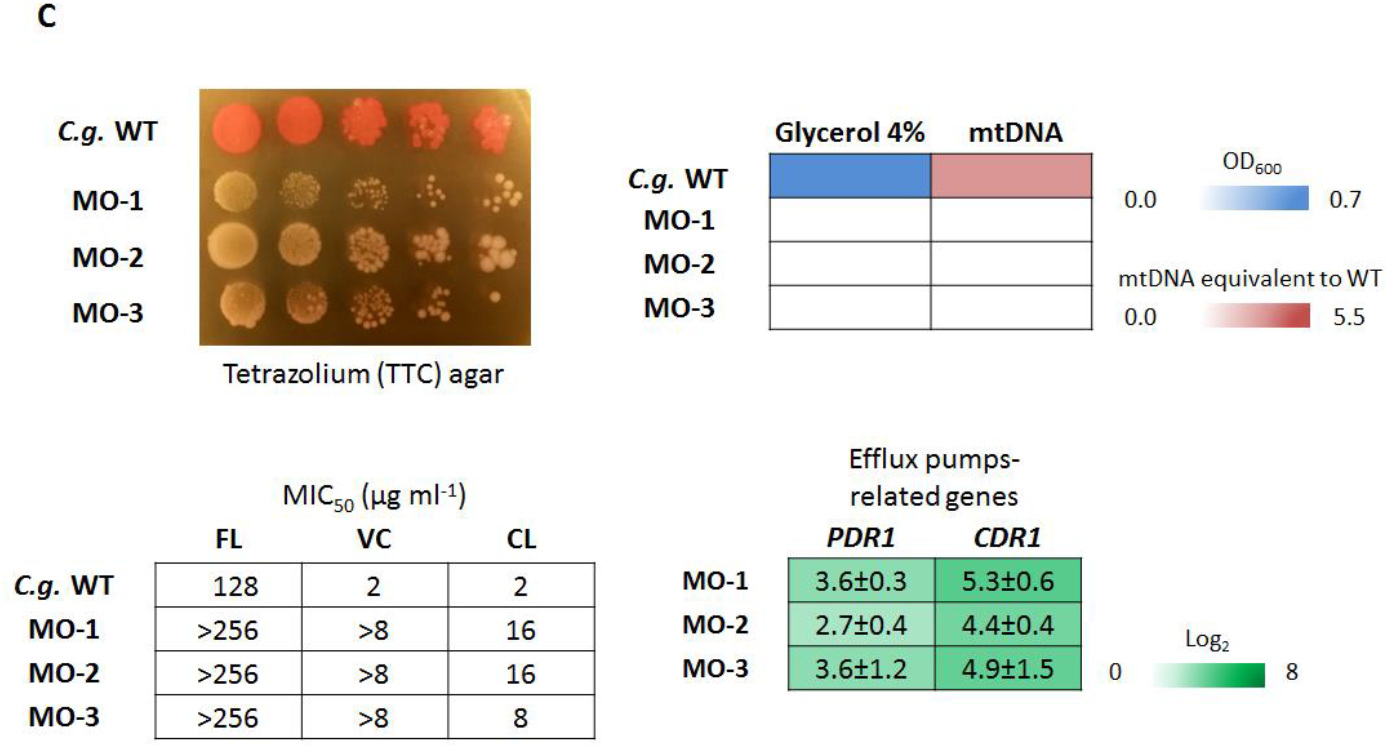
The *Petite* phenotype emerges from the wild type after phagocytosis. **(A)** *Cgmip1*Δ shows higher survival up to one day of co-incubation, but loses this advantage over extended periods of intracellular existence (n=5 with 1 different donor in each of the 5 independent experiments, each point represents a single survival test). **(B)** Cells with *petite* phenotypes arise from the wild type during co-incubation with hMDMs (red) at a much higher rate than the spontaneous appearance of *petite* in RPMI (blue). The time points with the highest frequency of *petite* emergence correspond to the time point of increased fitness of *Cgmip1*∆ during phagocytosis (Red: n=5 with 1 different donor in each of the 5 independent experiments and 4 technical replicates each. Blue: n=3 in 3 different experiments with 6 technical replicates. Mean ± SD of the frequencies of *petite* emergence at each time point). **(C)** Phenotype characterization reveals the hMDMs-derived strains (MO-1 to −3) to be indeed *petite*. From up-left: Small colony forming and lack of mitochondrial reductive power, lack of growth in alternative carbon sources, absence of mitochondrial (mt) DNA, high resistance to azoles (all n=3 with mean values or representative picture shown) and overexpression of efflux pumps-related genes (mean ± SD, n=3 independent experiments with 3 technical replicates each). FL: Fluconazole, VC: Voriconazole and CL: Clotrimazole. **(A)** Statistically significantly different values (unpaired, two-sided Student’s t-test on log-transformed ratios) are indicated by asterisks as follows: *, p ≤ 0.05; ***, p ≤ 0.001.

We analyzed several of the stable wild type-derived *petite* colonies and found – in the majority, but not all of them – a lack of detectable mtDNA. We selected three colonies from different experiments for further characterization of their *petite* phenotype (Figure 3). These lacked functional mitochondria as well as mtDNA and, importantly, also showed high azole resistance with constitutive expression of efflux-pumps related genes, similar to *Cgmip1*∆ (Figure 3).

We hypothesized that phagosomal ROS production may have contributed to the loss of mitochondrial function (Guo, Sun, Chen, & Zhang, 2013; Qin, Liu, Cao, Li, & Tian, 2011). We therefore incubated wild type yeasts in RPMI medium with sublethal concentration (10mM) of H_2_O_2_ for 24 hours and observed emergence of small colonies at a low frequency, which did not grow in glycerol (data not shown).

These results indicate that *petite* phenotype is adaptive within macrophages at early time points, but not at later times, probably due to the long period of starvation in the phagosome that prevents it to replicate. In agreement with this presumable advantage, *petite* phenotypes arise from the respiratory-competent yeasts after three hours to one day of phagocytosis, the same time period in which *Cgmip1*∆ shows a higher fitness. Importantly, we also found that the macrophage-induced *petites* can revert to a respiratory metabolisms when grows again in absence of stress.

### The *Petite* phenotype triggered by fluconazole increases survival of phagocytosis at early time points

It is known that azoles trigger (often temporary) mitochondrial dysfunction in *C. glabrata* (Kaur et al., 2004; Shingu-Vazquez & Traven, 2011) which leads to fluconazole resistance through the upregulation of the efflux pump genes *CDR1* and *CDR2*, with the former being especially important in this resistance (Brun et al., 2004; Sanglard et al., 2001; Zhang & Moye-Rowley, 2001). In fact, *petite* mutants have very occasionally been isolated from patients undergoing fluconazole treatment (Bouchara et al., 2000; Posteraro et al., 2006). Since our results showed an advantage of the genetically created *petite* strains after phagocytosis, we wondered whether fluconazole-induced *petites* share the same increased fitness. We therefore incubated wild type yeasts for 8 h in RPMI media with 8 µg/ml of fluconazole, half the concentration of the reported MIC_50_ for *C. glabrata* (“The European Committee on Antimicrobial Susceptibility Testing. Breakpoint tables for interpretation of MICs for antifungal agents.,” 2020). Again, we observed the appearance of small colonies with a *petite* phenotype (Figure 4). When these strains were co-incubated with macrophages for three and six hours, all fluconazole-induced *petites* showed better survival in macrophages at both time points (Figure 4).

**Figure 4.**
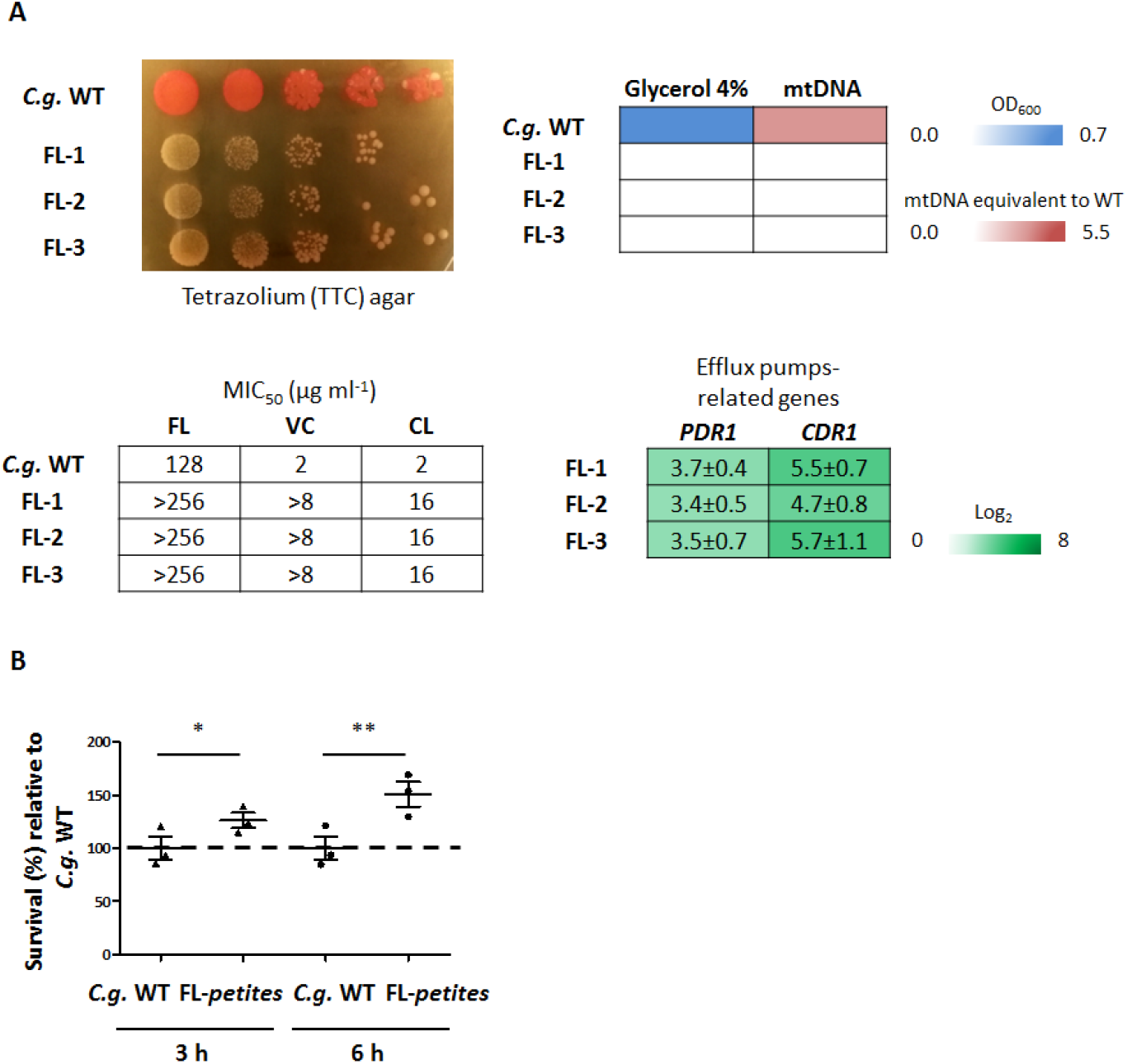
The *petite* phenotype triggered by fluconazole increases survival of phagocytosis at early time points. **(A)** Fluconazole-induced *petites* show *petite* phenotype similar to *Cgmip1*∆: Small colonies and lack of mitochondrial reductive power, lack of growth in alternative carbon sources, absence of mitochondrial (mt) DNA, high resistance to azoles (all n=3 with mean values or representative picture shown), and overexpression of efflux pumps-related genes (mean ± SD, n=3 independent experiments with 3 technical replicates each). FL: Fluconazole, VC: Voriconazole and CL: Clotrimazole. **(B)** Fluconazole-induced *petites* (FL-1 – FL-3) show better survival of phagocytosis at early time points (mean ± SD, n=3 with 1 donor in 3 independent experiments, each point represents a mean of 3 different colonies per donor, and each colony has 3 technical replicates). Statistically significantly different values (unpaired, two-sided Student’s t-test on log-transformed ratios) are indicated by asterisks as follows: *, p ≤ 0.05; **, p ≤ 0.01.

These results indicate a cross-resistance of the *petite* phenotype induced by and also protecting from both, phagocytosis and fluconazole: exposure to fluconazole triggers a higher fitness of *C. glabrata* inside macrophages and *vice versa*, fluconazole-resistant yeasts appear within macrophages.

### *Cgmip1*Δ shows higher basal expression of stress response-related genes and grows better under ER stress

To understand why switching to a *petite* phenotype increases intraphagosomal fitness, we measured the basal and induced expression of different stress-response genes and tested the resistance to infection-related stresses. *Cgmip1*∆ and the macrophages- and fluconazole-derived *petite* strains showed an increased basal expression of a range of environmental stress response genes (*HSP12, HSP42*, and *SGA1*) and cell wall stress-related genes encoding yapsins (*YPS1, YPS10*, and *YPS8*) (Figure 5). These genes have been shown to be highly up-regulated in the *petite* isolate BPY41, and may be involved in its hypervirulence (Ferrari, Sanguinetti, De Bernardis, et al., 2011). The yapsin *YPS* gene family has been implicated in the survival inside macrophages (Kaur, Ma, & Cormack, 2007), and we found that while the three *YPS* genes were up-regulated in *C. glabrata*, in the *S. cerevisiae petite* mutant (which does not survive phagocytosis better than its wild type) the most similar yapsin genes (*YPS1* and *MKC7*) showed no increased basal transcript levels (Figure S1).

**Figure 5.**
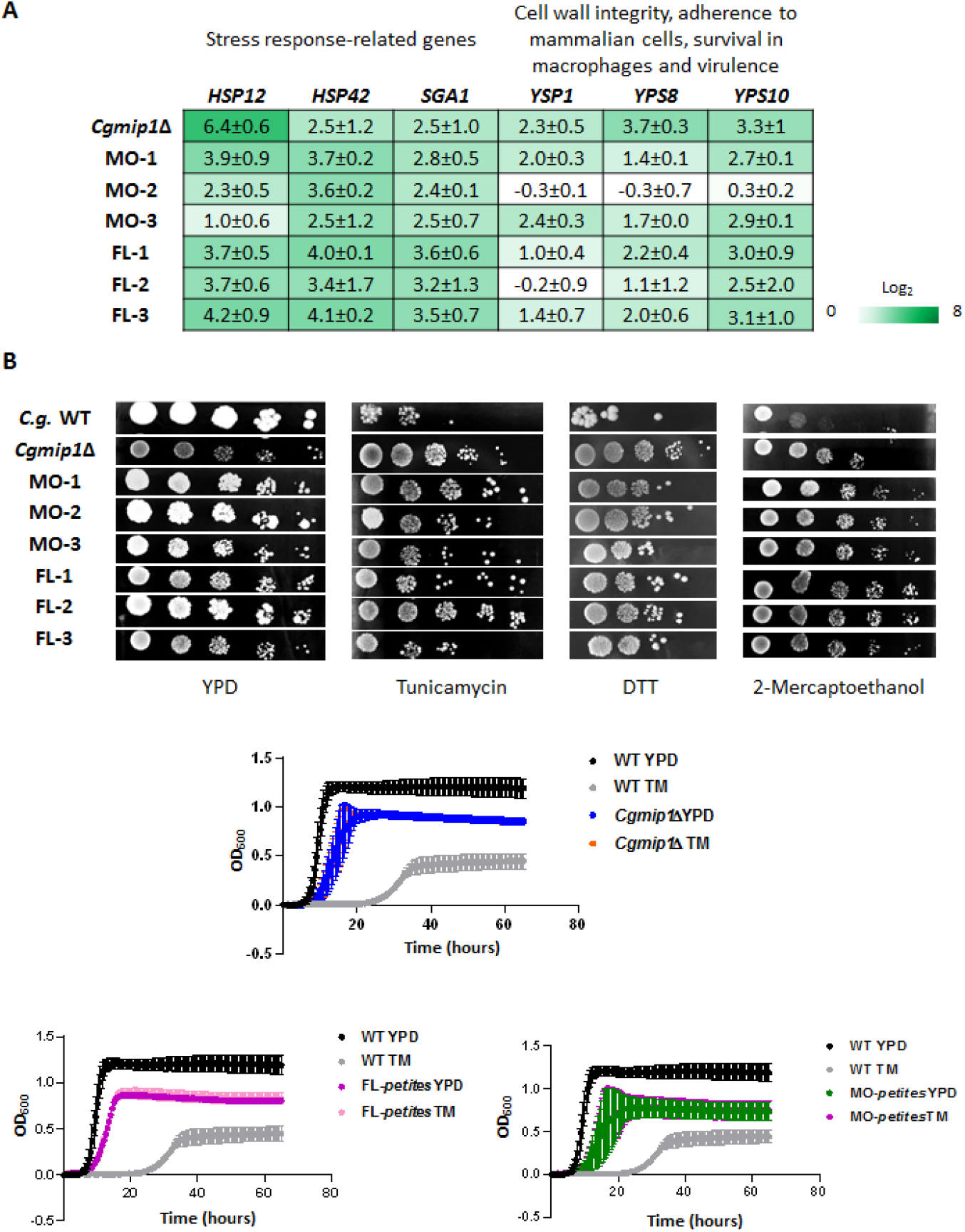
*Cgmip1*Δ shows higher basal expression of stress response-related genes and grows better under ER stress. **(A)** *Petite* variants of *C. glabrata* show high constitutive expression of stress-response genes even under non-stressed conditions (YPD) (mean ± SD, n=3 independent experiments with 3 technical replicates each), and **(B)** exhibit better growth than their wild type under different ER stresses on plates as well as with tunicamycin (TM) in liquid cultures (Mean ± SD, n=3 independent experiments or representative picture shown); FL-1 – FL-3: Fluconazole-induced *petites;* MO-1 – MO-3 macrophages-derived *petites*.

We then tested *Cgmip1*∆ in different *in vitro* stress conditions (data not shown). Interestingly, *Cgmip1*∆ showed oxidative stress resistance comparable to the wild type (Figure S1), in contrast to mitochondrial mutants of *S. cerevisiae* that are known to be especially sensitive to H_2_O_2_ (Thorpe, Fong, Alic, Higgins, & Dawes, 2004). Therefore we discarded a decreased sensitivity to oxidative stress as a possible explanation for the higher survival of *Cgmip1 ∆* within macrophages. However *Cgmip1*∆ and the azole- and macrophage-induced *C. glabrata petites* showed better growth than their wild type under ER stress conditions (Figure 5). Since efflux pumps play a central role in azole resistance, we analyzed mutants lacking their main transcriptional activator Pdr1 (Caudle et al., 2011; Thakur et al., 2008) in both, the wild type (*pdr1*Δ) and the *Cgmip1*∆ (*Cgmip1*Δ*+pdr1*Δ) background. As expected, both mutants grew poorly in increasing concentrations of fluconazole (Figure S1). However, under ER stress, the double mutant (lacking *CgMIP1* and *CgPDR1*Δ) still exhibited significantly better growth than the single mutant and the wild type (Figure S1), showing that the *Cgmip1*Δ resistance cannot be solely dependent upon Pdr1-regulated pathways or functions. Evidently, additional resistance mechanisms exist in the *petite* phenotype. In agreement with these results, the double mutant *Cgmip1*Δ*+pdr1*Δ showed significantly higher survival than the wild type after phagocytosis by hMDMs, whereas a *pdr1*Δ single mutant was actually killed more. Thus, the Pdr1 pathway seems to be involved in stress resistance and macrophage survival of these strains, but this cannot be the sole underlying mechanism (Figure S1). Overall, these results show that *petite* strains possess a constitutively active detoxifying response that, together with the overexpression of efflux pumps, confers ER stress resistance (in addition to azole resistance) and could explain the higher fitness within the phagosome observed at early time points.

**Figure S1.**
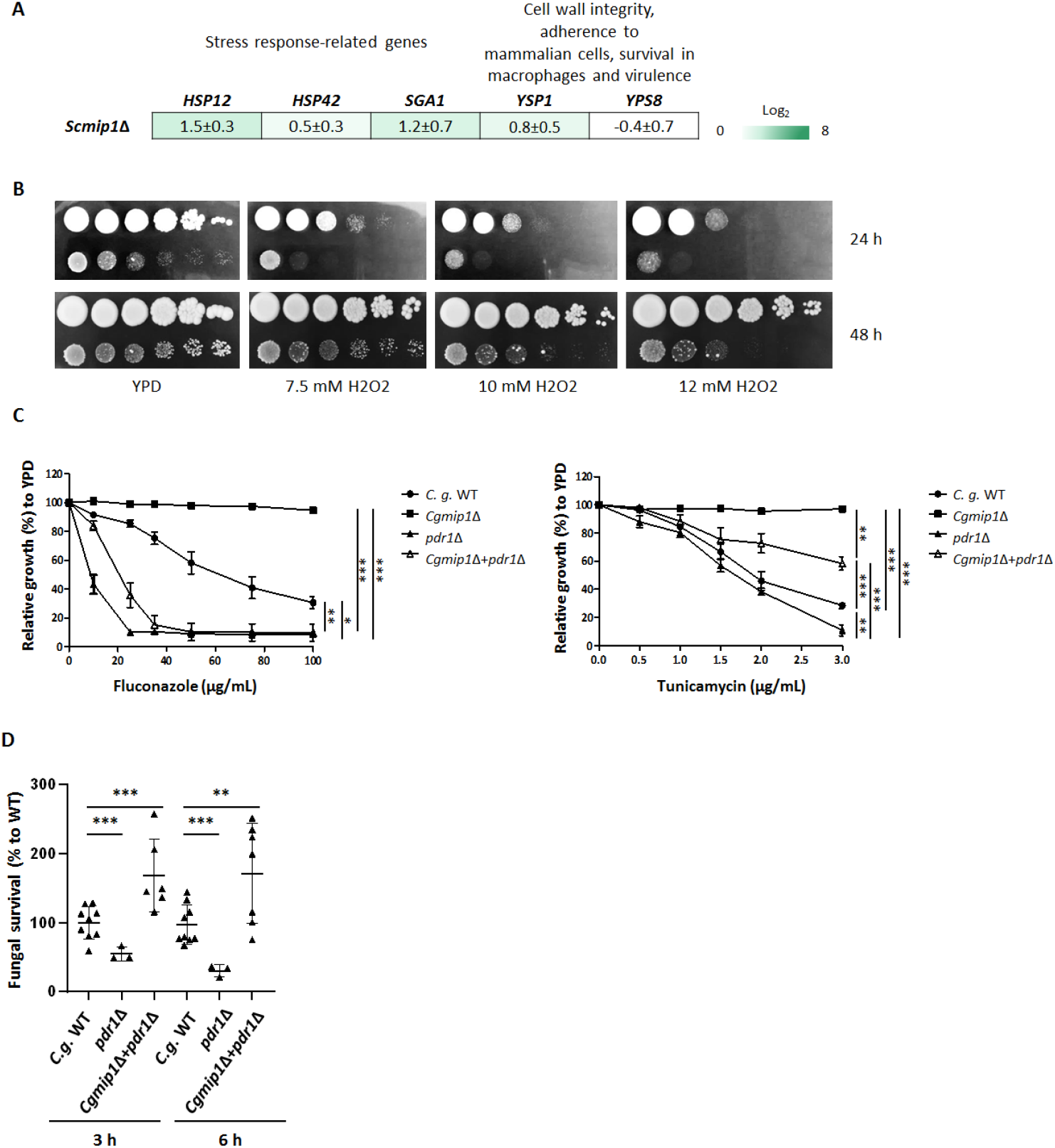
*YPS* genes and a *PDR1* pathway seem to be involved in the high survival to phagocytosis and ER stress resistance of *petite* phenotype. **(A)** *Scmip1*∆ does not show high constitutive expression of its orthologues of the *YPS* genes (*YPS1* and *MKC7* orthologue to *YPS8* in *C. glabrata*) (mean ± SD, n=3 independent experiments with 3 technical replicates each). **(B)** *Cgmip1*∆ shows a wild type-like level of resistance to oxidative stress (H_2_O_2_) on agar plates (representative picture shown). **(C)** The Pdr1 pathway seems to contribute to ER stress resistance but is not the solely responsible (mean ± SD, n=3 independent experiments with 3 technical replicates each) and **(D)** survival to phagocytosis by *pdr1*∆ single mutant and *Cgmip1*∆+*pdr1*∆ double mutant (mean ± SD, n=6 with 2 different donors in 3 independent experiments, each point represents the mean of 3 technical replicates). Statistically significantly different values (**C**: One-way ANOVA and Tukey’s test, **D**: unpaired two-sided Student’s t-test on log-transformed ratios) are indicated by asterisks as follows: *, p ≤ 0.05; **, p ≤ 0.01; ***, p ≤ 0.001.

### The *Cgmip1*∆ *petite* phenotype is adaptive under infection-like conditions but not in commensalism

We proposed that the generally reduced growth of *petite* mutants will be disadvantageous when competing with other microorganisms in a commensal environment, such as the human gut or vagina. Therefore, we wondered whether the emergence of *petite* phenotype may only be adaptive under infection-like conditions, such as antifungal treatment with azoles. To test this, we incubated separately or simultaneously wild type and *Cgmip1*Δ in the presence of vaginal cells and *Lactobacillus rhamnosus* for 24 hours mimicking a commensal-like situation (Graf et al., 2019), either in the absence or the presence of 8 µg/ml fluconazole. This fluconazole level is three times the concentration reported in vaginal fluids after a single oral dose (Houang, Chappatte, Byrne, Macrae, & Thorpe, 1990), but a concentration half lower than the MIC_50_ for *C. glabrata* (“;The European Committee on Antimicrobial Susceptibility Testing. Breakpoint tables for interpretation of MICs for antifungal agents.,” 2020). As expected, we observed only 60% of the wild type CFUs for the *petite* strain when it was incubated alone with human epithelial cells and without antifungals after 24 hours (Figure 6). The relative growth of *Cgmip1*Δ was further reduced in the presence of lactobacilli, and even more when co-incubated with the wild type and bacteria (Figure 6). However, the *petite* strain massively out-competed the wild type when fluconazole was present (Figure 6). Surprisingly, we did not observe a reduction in relative *Cgmip1*Δ CFUs in presence of both, lactobacilli and fluconazole, as it was seen when both species grew together in the absence of the drug.

**Figure 6.**
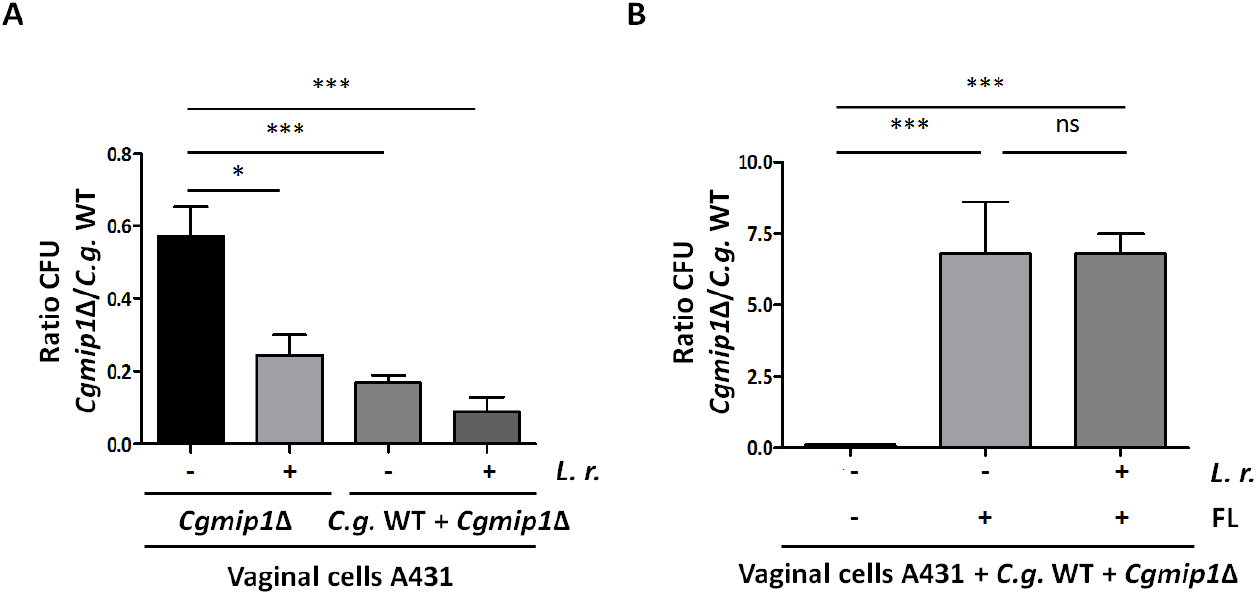
*Cgmip1*∆ *petite* phenotype is adaptive under infection-like conditions but not in commensal-like conditions. **(A)** The ratio of recovered *CgMIP1*-deleted to wild type *C. glabrata* cells is low when they were incubated separately on vaginal cells (*Cgmip1*Δ/-). The presence of *Lactobacillus rhamnosus* (*L. r*.) further shifts that ratio toward a lower recovery of *Cgmip1*Δ, and these effects are exacerbated in a direct competition in the same wells (*C. g*. WT + *Cgmip*Δ). Data shown is mean ± SD, n=3 independent experiments. **(B)** In an infection model in the presence of fluconazole (FL) the effect is inverted. Without fluconazole, *Cgmip*Δ is outcompeted as before (note the scale), but it has a decisive advantage in the presence of the antifungal drug, independent of the commensal bacteria (mean ± SD, n=3 independent experiments). **(A-B)** Statistically significantly different values (unpaired, two-sided Student’s t-test on log-transformed ratios) are indicated by asterisks as follows: *, p ≤ 0.05; ***, p ≤ 0.001.

In conclusion, these results show how in commensal-like and non-treated conditions *Cgmip1*Δ is outcompeted by respiratory-competent yeast cells and is also less able to compete with commensal bacteria. We conclude that this phenotype likely emerges only under conditions where it is advantageous, and then exists only transiently. These conditions include fluconazole treatment, but also uptake by macrophages.

### The *Cgmip1*Δ *petite* phenotype is found in clinical strains

Our data so far indicate that the *petite* phenotype should only appear transiently or at low rates in patients, but then provide significant advantages by increasing resistance to phagocytes and antifungals. We therefore screened two collections of 146 clinical *C. glabrata* isolates in total, provided by two different laboratories. Sixteen strains were identified as *petite*, i.e. they were showing a small colony size, absence of mitochondrial reductive power, and no growth in alternative carbon sources (Figure S2). The only common clinical characteristic these strains show is that majority of them was isolated from patients with underlying diseases (Table S1). These strains showed absence (but also sometimes even higher amounts) of mtDNA fragment we screened for. Furthermore, like our experimentally created *petites*, they exhibited high resistance to azoles, although they had not been exposed to azole treatment (Table S1), and grew better under ER stress compared to the wild type (Figure S2). Lastly, like the other *petite* variants, the clinical *petites* generally showed a higher survival inside of macrophages at early time points compared to the wild type (Figure S2). These results indicate that *petite* mutant can emerge during *C. glabrata* infections *in vivo* in clinical settings, and that these exhibit all the resistances we found in the experimentally created *petite* strains.

**Figure S2.**
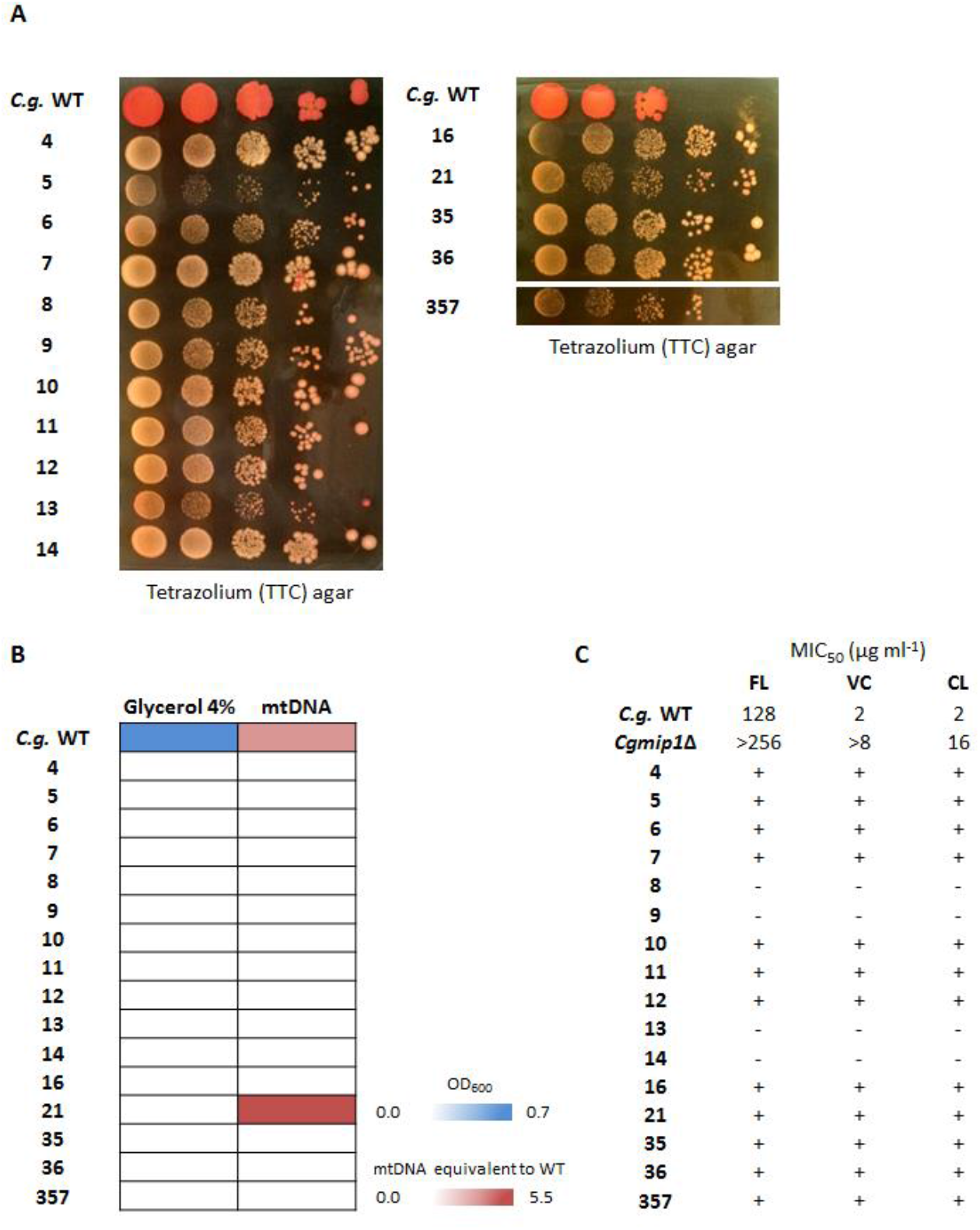
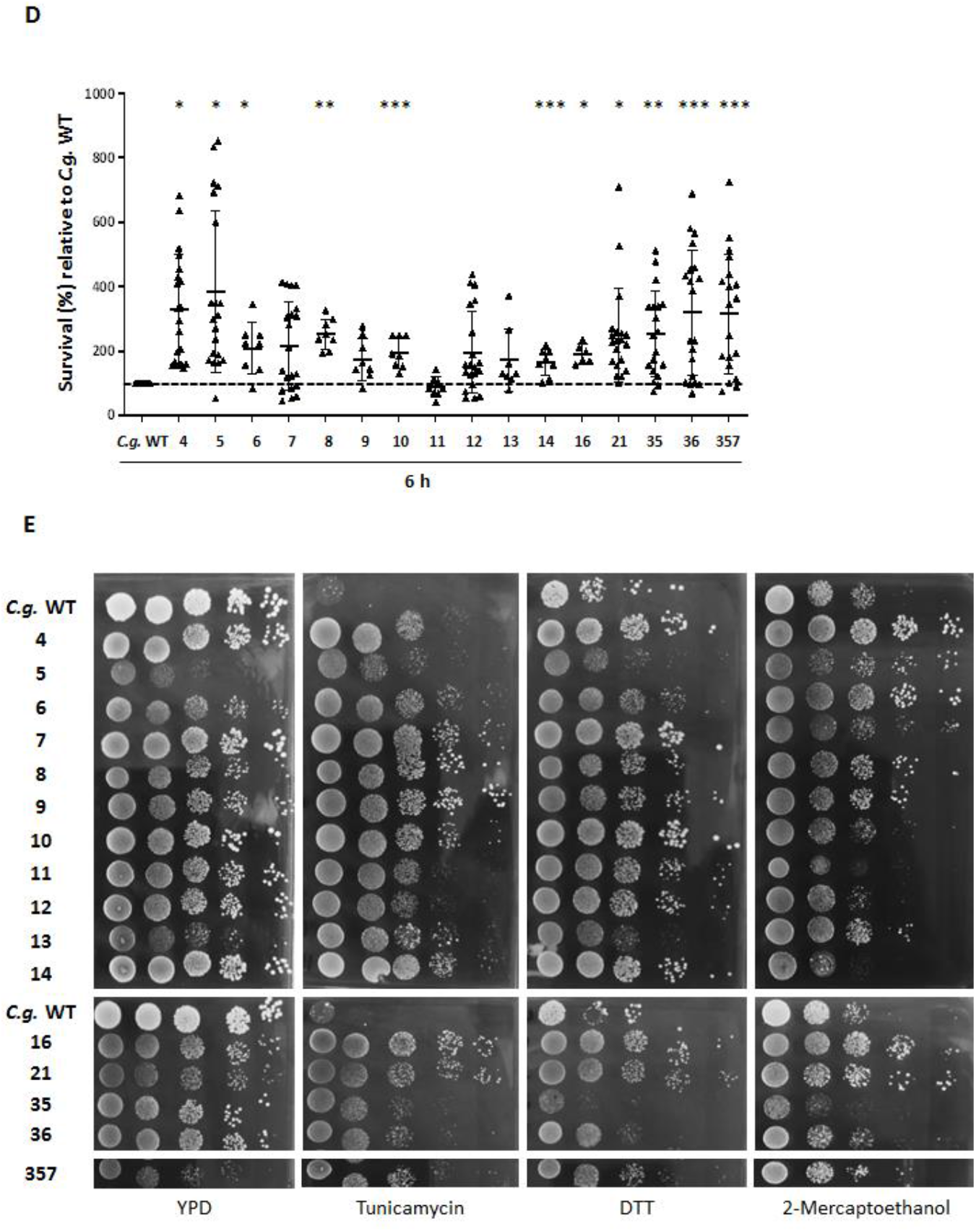
*Cgmip1Δ petite* phenotype is found in clinical strains. **(A)** The clinical *petite* variants show formation of small colonies and lack mitochondrial reductive power (representative picture shown). **(B)** None of them show growth in alternative carbon sources, and mostly absence of mitochondrial (mt) DNA (mean of n=3 independent experiments). **(C)** Most clinical isolates were able to grow (+) at the increased azole levels that indicated the resistance of the *petite* strains, which were generated *in vitro*. FL: Fluconazole, VC: Voriconazole and CL: Clotrimazole. **(D)** Clinical isolates with *petite* phenotype show increased survival to phagocytosis after 6 hours (mean ± SD, n=4 with 2 different donors in 2 independent experiments, each point represents a single survival test). Statistically significantly different values (One-way ANOVA and Dunnett’s test on log-transformed ratios) are indicated by asterisks as follows: *, p ≤ 0.05; **, p ≤ 0.01; ***, p ≤ 0.001. **(E)** *Petite* clinical strains generally grow better under ER stress than the ATCC 2001 reference strain (representative picture shown).

**Table S1.** Information about *petite* clinical strains of *C. glabrata* obtained from The Institute for Hygiene and Microbiology, Julius-Maximilians-University.

| Sample | Specimen | Sex | Age | Medical precondition | Disease | Immunosuppression | Antifungal therapy |
| --- | --- | --- | --- | --- | --- | --- | --- |
| 4 | urine sample | female | 50-55 | right heart failure with pulmonary hypertension | foreign body infection (atrial catheter with KNS) + pleural empyema (*E. cloacae*) | none | none |
| 5 |  |  |  |  |  |  |  |
| 6 | urine sample | female | 50-55 | Endometriosis | pyelonephritis with urosepsis | none | none |
| 7 | blood culture | female | 35-40 | Crohn's disease | sigmaperforation mit peritonitis, *C. glabrata* candidemia | biologicals, cortison high dose | caspofungin |
| 8 |  |  |  |  |  |  |  |
| 9 |  |  |  |  |  |  |  |
| 10 |  |  |  |  |  |  |  |
| 11 | expectorate | male | 30-35 | cystic fibrosis | Exacerbation | none | none |
| 12 |  |  |  |  |  |  |  |
| 13 |  |  |  |  |  |  |  |
| 14 | urine (bladder catheter) | female | 80-85 | cardiac bypass graft | compartment syndrome | none | none |
| 16 | punctate liverabscess | female | 75-80 | pancreatic adenocarcinoma with arterial erusion | abdominal abscess with liver necrosis | chemotherapy | caspofungin |
| 21 | abdominal punctate | male | 80-85 | colon adenocarcinoma, asthma b. | anastomotic leakage with adjunct peritonitis | none | none |
| 35 | sternocleidomastoid abscess membrane | male | 50-55 | ARDS (acute respiratory distress syndrome) due to COVID 19, chronic bronchitis | cervical abscess due to *C. glabrata* and catheter related mycosis | none | caspofungin |
| 36 |  |  |  |  |  |  |  |

### *CgMIP1* sequencing of *petite* clinical isolates

The fact that the mitochondrial polymerase gene *CgMIP1* shows a high value of positive selection (*d*_*N*_*/d*_*S*_=3.40) during the diversification of *C. glabrata* as a species and the presence of *petite* strains in clinical isolates of *C. glabrata* led us to hypothesize that these two phenomena are connected. We therefore searched for mutations of *CgMIP1* by obtaining the genome sequences of fourteen clinical strains isolated from seven different patients, which thirteen were *petite* and one respiratory competent. As comparison, we used the reference *C. glabrata* strain ATCC 2001 and sixteen respiratory-competent clinical strains, whose genome sequences have been previously obtained (Carrete et al., 2018). Comparing to the wild type strain, we found different mutations along the *MIP1* gene sequence, but we did not observe a specific common mutation pattern for the *petite* strains (Figure 7). Furthermore, none of these mutations were found in the predicted polymerase or exonuclease domains, which are important for the function of the protein (Lodi et al., 2015). Interestingly, we found variations of the N-terminal mitochondrial targeting sequence, which we determined by TargetP 2.0. Twelve *petites* (98.3%) show insertions of up to three more amino acids in the positions 24S, 25M and 26L/R, in comparison to the respiratory-competent strains, from which only six contained such insertions (35.3%). Furthermore, we looked for similar mutations in other proteins with known or expected mitochondrial localization. Dss1 is an exonuclease of the mitochondrial degradasome, and *CAGL0K03047g* is an ortholog of the *S. cerevisiae ABF2* gene, which has a role in mtDNA replication. Like Mip1, both are essential for mitochondria biogenesis (Razew et al., 2018) and maintenance (Diffley & Stillman, 1991). Hem1 is localized in the mitochondrial matrix and required for heme biosynthesis in *S. cerevisiae*. Pgs1 is a mitochondrially localized proteins whose deletion leads to increased azole resistance (Batova et al., 2008), and Pup1 is a mitochondrial protein that is upregulated by Pdr1 in azole-resistant strains (Ferrari, Sanguinetti, Torelli, Posteraro, & Sanglard, 2011). We did not detect variability similar to Mip1’s in any of these investigated protein sequence, independent on whether a well-defined mitochondrial transfer peptide was detectable (Abf2, Hem1) or not (Dss1, Pgs1, Pup1).

**Figure 7.**
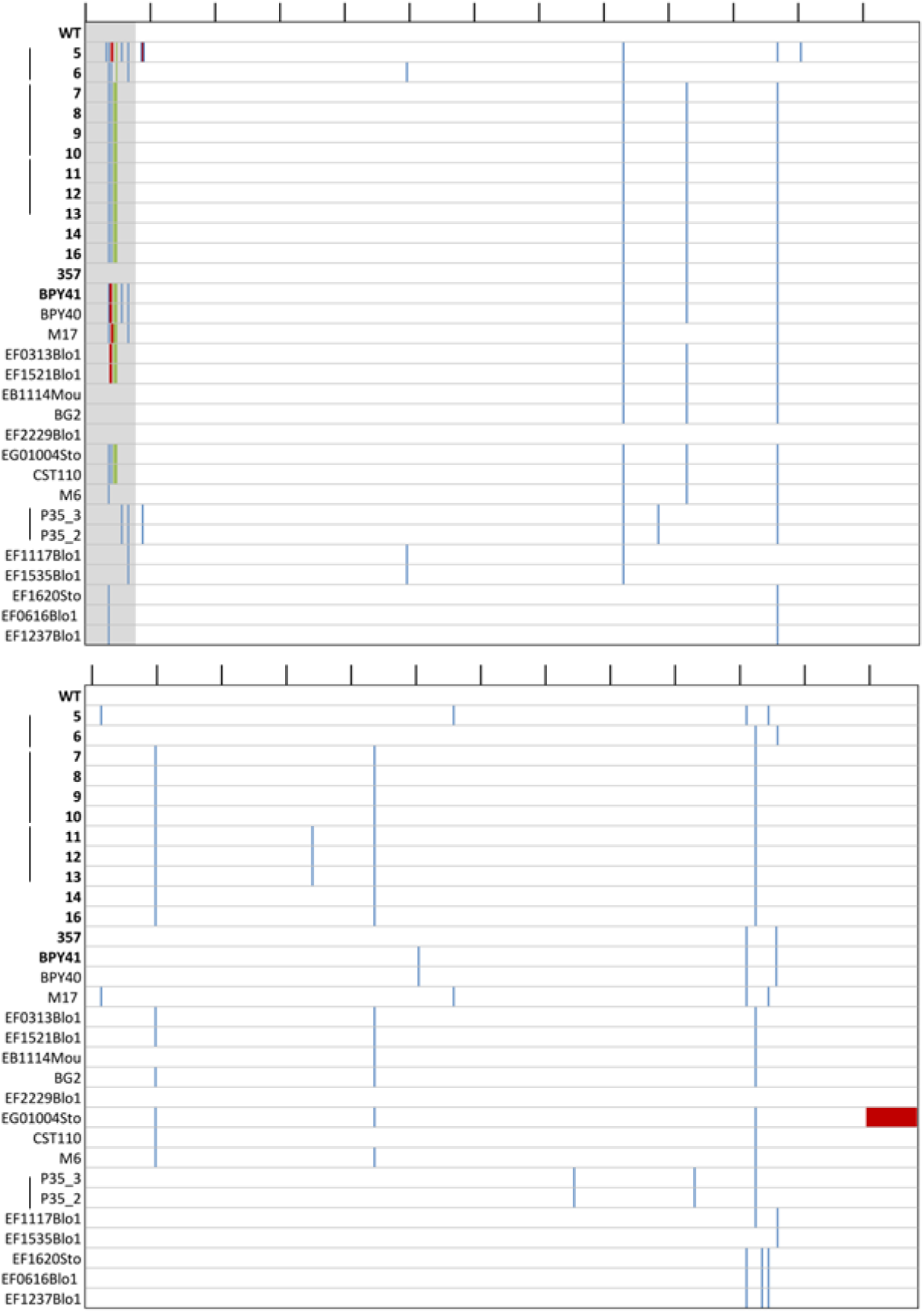
Mutation on the protein sequence of *CgMIP1* of *C. glabrata* clinical strains compared to the wild type. Wild type (ATCC 2001, WT), the first fourteen strains marked in bold are *petite* mutants, below there are the seventeen respiratory-competent strains. Grey: predicted mitochondrial signal peptide of the wild type, blue: Amino acid substitutions, green: Insertions, and red: Deletions. Black lines on top indicate the amino acid position every 50 amino acids. Black lines next to the strains’ names indicate that those strains have been isolated from the same patient.

## DISCUSSION

Despite the fact that *C. glabrata* is phylogenetically more closely related to the baker’s yeast *S. cerevisiae* than to the pathogenic *C. albicans*, it has by now become the most important non-*Candida albicans Candida* (NCAC) species to cause disease and is of growing concern in the clinics (Rodrigues, Silva, & Henriques, 2014). The relatively low host cell damage and immune responses that *C. glabrata* elicits, combined with its ability to survive and replicate within macrophages, indicates that stealth and persistence are its main virulence strategies during infection (Brunke & Hube, 2013). However, residing for a long time within the human body requires phenotypic plasticity to survive changing stresses, such as osmotic, ER, and oxidative stress, hypoxia, and starvation (Bliska & Casadevall, 2009; Casadevall, 2008; Gerwien, Skrahina, Kasper, Hube, & Brunke, 2018; Hube, 2009; Krishnan & Askew, 2014; Vylkova & Lorenz, 2014). One mechanism that leads to the better host adaptation and virulence in human pathogenic bacteria and fungi is the inactivation or loss of specific genes, which are then known as antivirulence genes (Bliven & Maurelli, 2012; Siscar-Lewin et al., 2019).

Specifically, in *C. albicans* and *C. glabrata* mutations decreasing mitochondrial function can affect host-pathogen interactions. Although *C. albicans* is considered a *petite*-negative species (Chen & Clark-Walker, 2000), a mutant with uncoupled oxidative phosphorylation was recovered after serial passaging of wild type *C. albicans* through murine spleens (Cheng, Clancy, Zhang, et al., 2007). It showed higher tolerance to ROS, altered cell wall composition, resistance to phagocytosis by neutrophils and macrophages, as well as an increased persistence and higher fungal burden in mice. Of note, in contrast to its progenitor, this strain did not kill infected mice any more. The authors pointed out that the uncoupled oxidative phosphorylation lowers intrinsic ROS production, which may be advantageous inside the phagosome, while the altered cell wall composition may diminish immune recognition. The mutant also showed a lower susceptibility to fluconazole and voriconazole due to increased expression of the efflux pump-encoding gene *MDR1* (Cheng, Clancy, Nguyen, Clapp, & Nguyen, 2007). In another example, a respiratory-competent *C. glabrata* strain and its *petite* mutant were sequentially isolated from a patient undergoing long-term fluconazole treatment (Ferrari, Sanguinetti, De Bernardis, et al., 2011; Posteraro et al., 2006). The mutant showed a high expression of virulence- and efflux pumps-related genes, oxido-reductive metabolism and stress response genes, as well as cell wall-related genes. It also led to a higher mortality in neutropenic mice and a higher tissue burden in immunocompetent mice (Ferrari, Sanguinetti, De Bernardis, et al., 2011), and it was also resistant to azoles due to the *PDR1*-dependent upregulation of the efflux pumps-encoding genes *CDR1* and *CDR2*. The mitochondrial-related gene, *PUP1*, was also strongly upregulated in a *PDR1*-dependent manner. Interestingly, enhanced virulence of *C. glabrata* associated with high upregulation of *CDR1* and *PUP1* has been observed in azole resistance clinical isolates resulting from gain of function mutations (GOF) in the *PDR1* gene (Ferrari, Sanguinetti, Torelli, et al., 2011). It was therefore speculated that both genes may contribute to favor *C. glabrata* in host interactions, in a still unknown manner. Thus, the *petite* phenotype may constitute a relevant pathogenic form of *C. glabrata*, and genes involved in mitochondrial function may be considered potential antivirulence genes (Siscar-Lewin et al., 2019).

In agreement with these previous findings, our study highlights the adaptive advantage that the lack of mitochondrial function has for *C. glabrata* under infection conditions. Likely not coincidentally, the mtDNA polymerase encoding gene *CgMIP1* seems to have been under selective pressures during the evolution of *C. glabrata* (Gabaldon et al., 2013). We found that deletion of this gene leads to loss of mtDNA and triggers the *petite* phenotype in both *C. glabrata* and *S. cerevisiae*. This phenotype confers a survival advantage at early time points after phagocytosis, but only for *C. glabrata* and not for *S. cerevisiae*. In addition, *petite* variants appeared from phagocytosed wild type cells at an appreciable frequency especially at early time points during their interaction with macrophages. We argue that these were likely induced by the intraphagosomal environment and provided an immediate advantage to the fungus. However, within the phagosome, glucose is absent and only alternative carbon sources are available (carboxylic acids, amino acids, peptides, N-acetylglucosamine, and fatty acids), which require mitochondrial oxido-reductive power for their metabolism (Lorenz et al., 2004; Sprenger et al., 2018). We assume that this is the reason that in the long term, after four and seven days, the advantage of the *petite* phenotype is reverted, as they starve and are recovered in ever lower numbers. Of note, the fact that in this experimental model oxygen is present at normal atmospheric levels may confer an advantage to the respiratory competent wild type. In infected tissue *in vivo*, in contrast, oxygen levels are low and it is known that hypoxia modulates innate immune response and enhance phagocytosis (Anand et al., 2007; Nizet & Johnson, 2009). In this case, the *petite* phenotype may even confer an adaptive advantage over respiratory-competent yeasts, which must rewire their metabolism upon confrontation with phagocytes. In addition, we observed that the *petite* phenotype can reverse if the fungi find themselves outside macrophages and the associated stresses. It seems feasible therefore that macrophage-induced *petites* regain their normal growth behavior *in vivo* if the fungus escapes the phagocytes and the change into a *petite* phenotype represents a temporary adaptation of *C. glabrata* to adverse conditions.

Importantly, our macrophage-derive*d petite* variants showed high resistance to azoles, in addition to other typical *petite* characteristics. In turn, fluconazole-derived *petites* showed the expected loss of susceptibility to azoles and surprisingly, a better survival to phagocytosis as well. It is known that fluconazole can trigger (temporary) loss of mitochondrial function in *C. glabrata* and *S. cerevisiae*, and as a consequence an increased fluconazole resistance (Brun et al., 2004; Kaur et al., 2004; Sanglard et al., 2001; Zhang & Moye-Rowley, 2001). However, to our knowledge this study shows for the first time that phagocytosis and intraphagosomal residence can lead to the emergence of fluconazole resistance, and *vice versa*, in a potentially clinically important cross-resistance phenomenon.

The mechanistic basis of how these resistances develop is not completely understood: the clinical *petite* variants reported so far have all been isolated from patients undergoing fluconazole treatment (Bouchara et al., 2000; Posteraro et al., 2006) and most of the *petite* isolates analyzed here lack mtDNA, but synthesis inhibition or degradation of mtDNA by the action of azoles has not yet been reported. In fact, it was shown that azole exposure does not always lead to a loss of mtDNA, but rather damages mitochondrial components (Kaur et al., 2004; Sanglard et al., 2001; Zhang & Moye-Rowley, 2001). Fluconazole-induced *petite* can revert at a frequency of 1.5 × 10^−2^ (Kaur et al., 2004), which suggests a genetic or epigenetic regulation. Our data indicates that *C. glabrata* turns *petite* within the macrophages due to phagosomal oxidative stress, since we observed similar conversions during incubation with H_2_O_2_. It is known that oxidative stress can trigger mitochondrial damage and loss of activity by affecting mitochondrial membrane permeability, the respiratory chain or the mtDNA (Guo et al., 2013; Qin et al., 2011). These factors could be at work during the induction of the *petite* phenotype in macrophages. Importantly, like azole-induced *petites*, we found that a fraction of these macrophage-derived *petite* phenotypes were reversible.

The loss of mitochondrial function also activates compensatory pathways, such as detoxifying mechanisms and cell wall remodeling (Shingu-Vazquez & Traven, 2011). Indeed, all our *petite* strains showed constitutive expression of efflux pumps-related (and azole resistance-mediating) genes *PDR1* and *CDR1*, and some transcription of heat shock protein-encoding genes like *HSP12* and *HS42*, and *SGA1* (Ferrari, Sanguinetti, De Bernardis, et al., 2011; Moskvina, Schuller, Maurer, Mager, & Ruis, 1998). Moreover, all *C. glabrata petites* exhibited a high resistance to ER stressors. We suggest that this ER stress resistance is acquired via mitochondria dysfunction and confers an adaptive advantage in the phagosome, since it has been shown that ROS may not act alone in killing the fungus but in combination with additional stresses: Suppression of ROS by macrophages alone does not increase the fungal survival (Kasper et al., 2015). In addition this ER stress resistance would also confer protection against the generic ER stress that pathogens face during infection (Cohen, Lobritz, & Collins, 2013; Kaur et al., 2007; Tiwari, Thakur, & Shankar, 2015). We also showed that the *PDR1* pathway is at least partially involved in the ER stress resistance as well as in macrophages survival, in agreement with previous findings (Ferrari, Sanguinetti, Torelli, et al., 2011). The majority of *petites* also showed increased expression of members of the *YPS* gene family. It has been suggested that Yps-mediated cell wall remodeling can play a role in altering or suppressing macrophage activation (Kaur et al., 2007), and we hypothesize that this may contribute to the better survival of *petite* variants within the phagosome. Of note, *S. cerevisiae*, which did not benefit from a *petite* phenotype in macrophages, also did not show such a constitutive *YPS* orthologue expression. In addition, it has been reported that different mitochondrial mutants of *S. cerevisiae* show increased sensitivity to oxidative stress by H_2_O_2_ (Thorpe et al., 2004), which was not the case for our *Cgmip1*Δ mutant, and this phenomenon should therefore not influence its survival in macrophages. The altered cell wall, and especially the increased mannan exposure of *Cgmip1*Δ, can explain the increased phagocytosis rate, as this is known to be mannan-dependent (Keppler-Ross et al., 2010; Snarr, Qureshi, & Sheppard, 2017). Clearly, the induction of a *petite* phenotype by either azoles or phagocytosis had a strong influence on the macrophage-fungus interactions, benefitting *C. glabrata* in the phagosome.

The *petite* phenotype shows nonetheless a strong handicap in fitness and competitiveness in our commensal-like model, likely due to its slow growth, as observed before (Ben-Ami & Kontoyiannis, 2012). However, in our model of vaginal candidiasis treated with fluconazole, the *petite* phenotype shows a steady advantage. Therefore, we suggest that the *petite* phenotype, which also appears naturally and in absence of stress at low frequencies, serves as a bet-hedging strategy to face stressful conditions, such as phagocytosis or azole exposure, in *C. glabrata*. Similar phenotypes are described in bacteria, for which it is known that microorganisms that give rise to heterogeneous populations and phenotypic switching are more likely to survive in fluctuating environments, than otherwise “stable” populations (Arnoldini et al., 2014; Holland, Reader, Dyer, & Avery, 2014). Important intracellular pathogens like *Staphylococcus aureus* and *Salmonella* are known to form small colony variants (SCVs), an analogous phenotype of *petite*, as part of the bacterial heterogeneity that might confer an adaptive advantage upon environmental changes (Arnoldini et al., 2014; Day, 2013; Tuchscherr et al., 2019). These show a decrease in antibiotic susceptibility, link to chronic and relapsing, often therapy-refractory infections. Moreover, they show reduced expression of virulence factors, and higher adhesion, which promotes internalization in host cells and facilitate immune-evasion and long-term persistence within their hosts (Kahl, Becker, & Loffler, 2016; Proctor et al., 2006). SCVs from many gram-positive and -negative bacteria have been recovered from clinical tissues (Kahl et al., 2016; Proctor et al., 2006). On the yeast counterpart, so far only a *petite* mutant of *C. glabrata* has been reported to possess any pathogenic advantage (Ferrari, Sanguinetti, De Bernardis, et al., 2011). Analogously to the SCVs, these mutants have been isolated from cases of antifungal treatment (Bouchara et al., 2000) and recurrent fungemia (Posteraro et al., 2006), with a decreased susceptibility to antifungals and increased fungal burden in animal models of infection (Ferrari, Sanguinetti, De Bernardis, et al., 2011). Furthermore, recently it has been shown that a negative correlation between fitness costs derived from drug resistance and virulence is not always the case, but in fact virulence could be maintained or even increased in the presence of such costs that results in a reduced growth rate in *C. glabrata* (Duxbury, Bates, Beardmore, & Gudelj, 2020). This agrees with our hypothesis of *petite* phenotype as an advantageous strategy during infection despite its slower growth.

Given the advantages of *petite* variants during infection, one would expect to frequently find *C. glabrata petite* phenotypes in clinical samples; but although *petites* have been reported (Bouchara et al., 2000; Posteraro et al., 2006), these reports seem to be rare. Nonetheless, when we specifically looked for *petite* phenotype in clinical isolates we found seventeen of these strains. They shared all the hallmark features with *Cgmip1*∆, macrophage-, and fluconazole-derived *petites*. The majority showed loss of mtDNA, azole resistance, and increased survival upon phagocytosis. It may well be that *C. glabrata petite* phenotypes are actually more common in clinical samples, but potentially overlooked because of their slow growth.

We started these investigations because of the signs of recent selection on the mitochondrial DNA polymerase gene *CgMIP1*. We hypothesized that this could indicate a role in the adaptation of *C. glabrata* to host environments. Indeed, some (but not all) of the *petite* clinical isolates we investigated showed mutations in the *CgMIP1* gene sequence and the majority showed a insertion pattern in the presumable mitochondria targeting sequence, which may relate to their *petite* phenotypes via a reduction of mitochondrial polymerase function. In addition, we found that other mitochondria-related proteins did not show similar mutation frequency, which supports the idea that CgMip1 may still be a target for selection. It has been shown that point mutations in the polymerase domain of the orthologous *MIP1* gene in *S. cerevisiae* triggered emergence of *petites*, and this frequency increased with higher temperatures. Specifically, the mutation E900G yielded from 6% *petites* at 28°C to 92% at 36°C (Baruffini, Ferrero, & Foury, 2007). While many of these petites were rho^0^ – completely devoid of mtDNA – some still contained amplified mtDNA fragments that map to various positions of the mtDNA and were considered rho^−^ (Baruffini et al., 2007). These mtDNA fragments can rescue strains that are respiratory deficient due to mutations in mtDNA by recombination after crossing (Baruffini et al., 2006; Tzagoloff, Akai, Needleman, & Zulch, 1975). Interestingly, we also found mtDNA fragments in our sequence data of clinical petite strains (data not shown), similar to *S. cerevisiae* rho^−^. Moreover, we also found that wild type strain of *C. glabrat*a and its two most closely related human pathogens, *Nakaseomyces nivariensis* and *Nakaseomyces bracarensis*, show the substitution E926D in Mip1 when compared to the environmental species *Nakaseomyces delphensis, Nakaseomyces bacillisporus*, and *Candida castelli*. Since this is equivalent to E900 position in *S. cerevisiae*, which upon mutation increases the frequency of *petite* occurences, we hypothesize that on the one hand *C. glabrata* could more easily turn *petite* than environmental *Nakaseomyces* species, but on the other hand it can also retain mtDNA in its genome as a possible mechanism to recover mitochondrial function once the *petite* phenotype is not adaptive. These tantalizing hypotheses will be tested in the near future.

Alternatively, the *C. glabrata CgMIP1* gene sequence may be the target of epigenetic regulation, as epigenetics have been suggested to be involved in the reversion of *petite* phenotype to wild type (Kaur et al., 2004). In these models, mtDNA levels are reduced down to a (near) complete loss of mitochondrial function. Upon resumption of polymerase function, the mitochondria function and growth then reverts to normal. The ability to switch between *petite* and non-*petite* phenotype would confer *C. glabrata* an important phenotypic plasticity to adapt to the host’s changing environments: The *petite* phenotype is less competitive as a commensal, but much fitter in infection situations with active phagocytes and antifungal treatment.

In conclusion, this study shows how temporary mitochondria dysfunction triggers a *petite* phenotype in *C. glabrata* under infection-like conditions, with the potential to confer cross-resistance between the macrophages and azole antifungal treatment. This has three implications. First, it adds mitochondrial function to the list of potential antivirulence factors, since its loss results in a gain in virulence potential. Second, it has implications for the treatment of *C. glabrata*, as fluconazole may inadvertently increase the fitness of the fungus against the innate host defenses. Due to the *petite* morphology and long generation times, such resistant isolates may then be missed in standard diagnostics. Third, our observations provide a potential clinical relevant route for the emergence of azole resistance of *C. glabrata* by immune activities, a new paradigm in the development of antifungal resistance.

## MATERIALS AND METHODS

### Screening and acquisition of suspected petite *C. glabrata* strains

In the course of routine diagnostics – executed by the Institute of Hygiene and Microbiology in Würzburg – all accumulated chromogenic candida agar plates (CHROMagar, Becton Dickinson, New Jersey, USA) were systematically collected and incubated for at least 7 days at 37°C prior to screening. After incubation plates were visually screened concerning growth, color and morphology. Only suspected *C. glabrata* small colony variants were subisolated and reincubated for a minimum of 3 days at 37°C. In case of a confirmed growth behavior final species identification was executed by MALDI-TOF (BioMerieux, Paris, France).

466 from a total of 3756 agar plates – originating from various clinical specimen examined between November 2019 and June 2020– exhibited growth after incubation. 525 different strains were identified through chromogenic media, of which the majority (312) presented itself in a green color suggesting *C. albicans*, whereas in 170 cases mauve colonies were observed. 82 of these were identified as *C. glabrata* using MALDI-TOF. Based on morphology, 41 strains were suspected to be *petites*, which was finally confirmed in 20 cases.

### Strains and growth conditions

All strains used in this study are listed in Table S2. *C. glabrata* mutant strains are derivatives of the laboratory strain ATCC 2001. In each strain a single open reading frame (ORF) was replaced with a bar-coded NAT1 resistance cassette in the strain ATCC 2001. All yeast strains were routinely grown overnight in YPD (1% yeast extract, 1% peptone, 2% glucose) at 37°C in a 180 rpm shaking incubator.

**Table S2.**
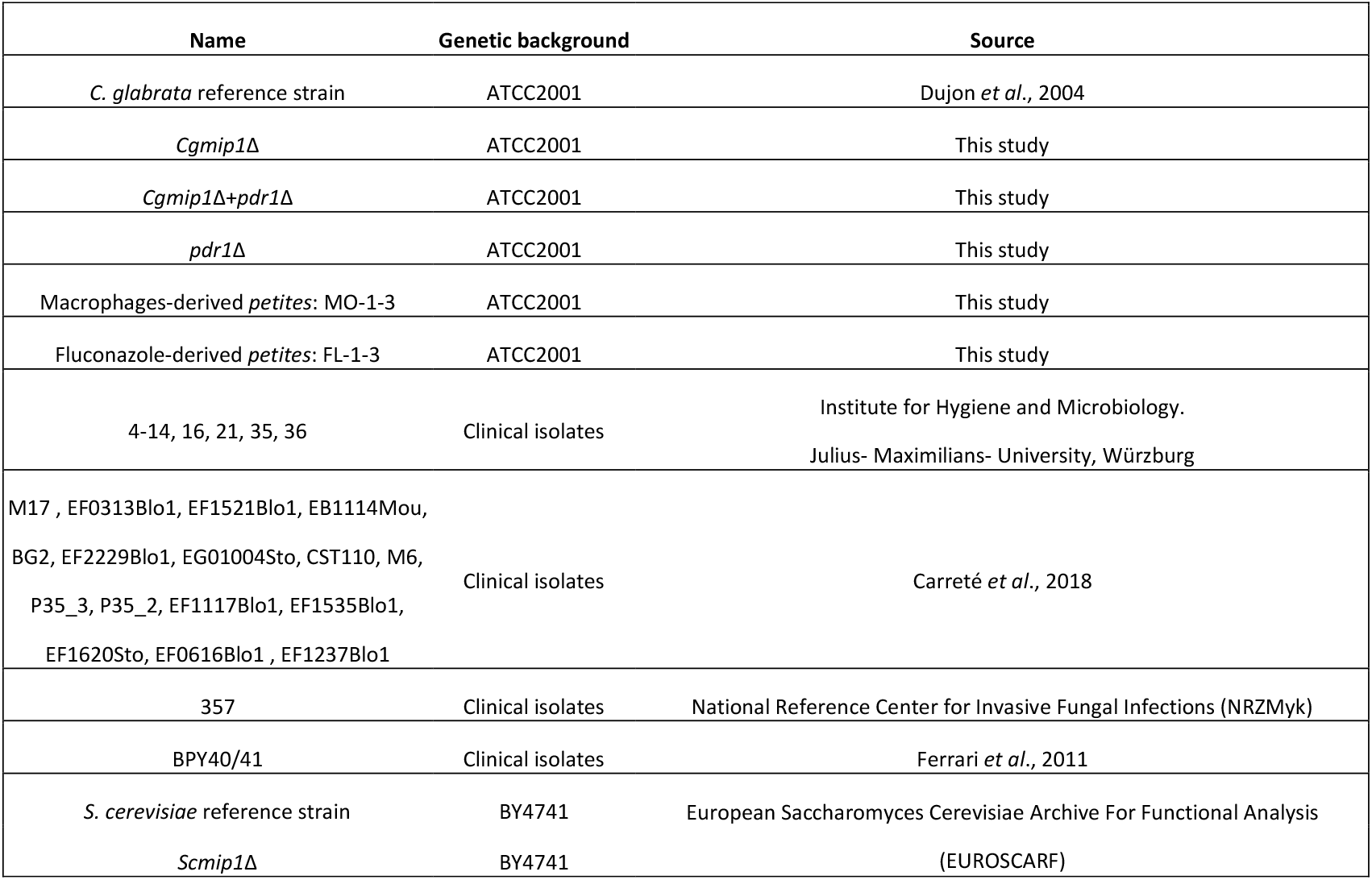
Strains used in this study.

To analyze sensitivity to H_2_O_2_ and ER stress, 5 µl of serial diluted yeast cultures (10^7^, 10^6^, 10^5^, 10^4^, 10^3^, 10^2^ cells/mL) were dropped on solid YPD media (YPD, agarose 2%) containing increasing concentrations of H_2_O_2_ (7.5 mM, 10 mM and 12 mM), Tunicamycin (2 µg/ml), DTT (10 mM) or 2-Mercaptoethanol (12 mM). Pictures were taken after 24 and 48 hours of incubation at 37°C. Mitochondria activity was visualized by growing serial dilutions of the strains on YPD agar containing 0.02% TTC (2,3,5-Triphenyltetrazolium chloride)(Sigma-Aldrich), and incubating cells in minimum media (1% yeast nitrogen base, 1% amino acids, 0.5% ammonium sulfate) with 4% glycerol as sole carbon source for 4 days at 37°C in a 180 rpm shaking incubator.

### Growth assays

To analyze stress sensitivity, five microliters of a yeast cell suspension (2×10^7^ cells/mL) was added to 195 µL media in a 96-well plate (Tissue Culture Test Plate, TPP Techno Plastic Products AG) containing liquid YPD or YPD and different concentrations of fluconazole (10, 25, 35, 50, 75, 100 µg/ml) and tunicamycin (0.5, 1, 1.5, 2 and 3 µg/ml). For the ER stress analysis yeast were incubated in YPD and tunicamycin (1.5 µg/ml). The growth was monitored by measuring the absorbance at 600 nm every 30 min for 150 cycles at 37 °C, using a Tecan Reader (Plate Reader infinite M200 PRO, Tecan Group GmbH) with orbital shaking. All experiments were done in independent biological triplicates on different days, and shown as mean with standard deviation (SD) for each time point.

The Minimum Inhibitory Concentration (MIC_50_) was done in a 96-well plate (Tissue Culture Test Plate, TPP Techno Plastic Products AG) containing minimum media (1% yeast nitrogen base, 1% amino acids, 0.5% amonio sulfate and 2% glucose) with increasing concentrations of fluconazole (FL) (4, 8, 16, 32, 64, 128, 256 µg/ml), voriconazole (VC) (0.5, 1, 2, 4, 8, 16 µg/ml) and clotrimazole (CL) (1, 2, 4, 8, 16 µg/ml) at 37°C.

### Fungal RNA isolation

For preparation of RNA from *in vitro* cultured *Candida* cells, stationary phase yeast cells were washed in PBS and OD was adjusted to 0.4 in 5 mL liquid YPD. After 3 h, cells were harvested and centrifuged. The isolation of the fungal RNA was performed as previously described (Lüttich, Brunke, & Hube, 2012). The RNA was then precipitated by adding 1 volume of isopropyl alcohol and one tenth volume of sodium acetate (pH 5.5). The quantity of the RNA was determined using the NanoDrop Spectrophotometer ND-1000 (NRW International GmbH).

### Expression analysis by reverse transcription-quantitative PCR (qRT-PCR)

cDNA was synthesized from DNase-treated RNA (1000 ng) using 0.5 µg oligo-dT_12-18_, 100 U Superscript^™^ III Reverse Transcriptase and 20 U RNaseOUT^™^ Recombinant RNase Inhibitor (all: Thermo Fischer Scientific) in a total volume of 20 µL for 2 h at 42 °C followed by heat-inactivation for 15 min at 70 °C. Quantitative PCR with EvaGreen^®^ QPCR Mix II (Bio&SELL) was performed with 1:10 diluted cDNA. Primers (Table S3) were used at a final concentration of 500 nM. Target gene expression was calculated using the ∆∆Ct method (Pfaffl, 2001), with normalization to the housekeeping genes *CgACT1* for *C. glabrata* or *ScACT1* for *S. cerevisiae*. For mtDNA quantification, yeast DNA was extracted following Harju *et al*., protocol (Harju, Fedosyuk, & Peterson, 2004) and 100 ng was the reaction concentration. *ScCOX3* and *CgCOX3* were used as a mitochondrial target gene and *CgACT1* or *ScACT1* as housekeeping genes. All experiments were done in independent biological triplicates on different days, and shown as mean with standard deviation (SD) for each time point.

### Chitin, mannan and β-glucan exposure

To measure chitin content, yeasts from an overnight culture were washed in PBS and incubated with 9 ng/ml of WGA-FITC diluted in PBS for 1 h at room temperature. After washing with PBS, fluorescence was quantified by flow cytometry (BD FACS Verse^®.^ BD Biosciences, Franklin Lakes, USA) counting 10,000 events. For mannan quantification, yeast cells were washed with PBS and incubated with concanavalin A-647 for 30 min at @37°C. After washing with PBS, fluorescence was quantified also by flow cytometry. For β-glucan staining, yeast cells were washed with PBS and incubated with 2% bovine serum albumin (BSA) for 30 min at 37°C, followed by a first step of 1 h of incubation with a monoclonal anti-β-glucan antibody (Biosupplies) (diluted 1:400 in 2% BSA) and a second step of 1 h of incubation with an Alexa Fluor 488 conjugate secondary antibody (Molecular Probes) (diluted 1:1000 in 2% BSA). Fluorescence was again quantified by flow cytometry. All experiments were done in independent biological triplicates on different days, and shown as mean with standard deviation (SD) for each time point.

### Isolation and differentiation of human monocyte-derived macrophages (hMDMs)

Blood was obtained from healthy human volunteers with written informed consent according to the declaration of Helsinki. The blood donation protocol and use of blood for this study were approved by the Jena institutional ethics committee (Ethik-Kommission des Universitätsklinikums Jena, Permission No 2207–01/08). Human peripheral blood mononuclear cells (PBMCs) from buffy coats donated by healthy volunteers were separated through Lymphocytes Separation Media (Capricorn Scientific) in Leucosep™ tubes (Greiner Bio-One) by density centrifugation. Magnetically labeled CD14 positive monocytes were selected by automated cell sorting (autoMACs; MiltenyiBiotec). To differentiate PBMC into human monocyte-derived macrophages (hMDMs), 1.7×10^7^ cells were seeded into 175 cm^2^ cell culture flasks in RPMI1640 media with L-glutamine (Thermo Fisher Scientific) containing 10 % heat-inactivated fetal bovine serum (FBS; Bio&SELL) and 50 ng/mL recombinant human macrophage colony-stimulating factor M-CSF (ImmunoTools). Cells were incubated for five days at 37 °C and 5 % CO_2_ until the medium was exchanged. After another two days, adherent hMDMs were detached with 50 mM EDTA in PBS and seeded in 96-well plates (4×10^4^ hMDMs/well) for survival-assay, in 12-well-plates (4×10^5^ hMDMs/well) for intracellular replication assay with 100 U/ml γ-INF, and 24-well-plates for long-term experiment (1.5 ×10^5^ hMDMs/well) without γ-INF. Prior to macrophage infection, medium was exchanged to serum free-RPMI medium and 100 U/ml γ-INF. For long-term experiment, medium was exchanged to RPMI1640 containing 10 % human serum (Bio&Sell 1: B&S Humanserum sterilised AB Male, Lot: BS.15472.5).

### Phagocytosis survival assay

Mutant strains were washed in phosphate buffered saline (PBS), and total numbers of cells were assessed by the use of a hemocytometer. MDMs in 96-well-plates were infected at an MOI of 1, and after 3 h and 6 h of coincubation at 37°C and 5% CO_2_, non-cell-associated yeasts were removed by washing with RPMI 1640. To measure yeast survival in MDMs, lysates of infected MDMs were plated on YPD plates to determine CFU.

The long-term experiment was perform in 24-well-plates were the cells were infected with a MOI of 1 and incubated for one week at 37°C and 5% CO_2_. After 3 h of coincubation, cells were washed with PBS and medium was exchanged to RPMI 1640 containing 10 % human serum. For 3 h time point both supernatant and lysate were plated. Until 1 day and 7 days times points, third of the medium was exchanged every day with in RPMI 1640 with 10% human serum. Then, only the lysate was plated. The lysate of 4 different wells was diluted accordingly and 200 CFU were plated on 4 YPD agar plates, which were afterwards incubated for 48 h at 37°C. The frequency of *petites* was determined via small colonies that were unable to grow with 4% glycerol as a sole carbon source in minimal medium (1% yeast nitrogen base, 1% amino acids, 0.5% ammonium sulfate). The growth was assayed for 3 days at 37°C in a 180 rpm shaking incubator. The frequency of spontaneous *petites* was calculated by incubation of 1.5×10^5^ cells ml^-1^ in RPMI 1640 for 7 days. At 3 hours, 1 day, 4 days, and 7 days samples were collected, diluted and 200 CFU were plated on 6 YPD agar plates, which were incubated for 48 h at 37°C.

### Replication within hMDMs

To quantify yeast intracellular replication, *C. glabrata* cells were labeled with 0.2 mg/mL fluorescein isothiocyanate (FITC) (Sigma-Aldrich) in carbonate buffer (0.15 M NaCl, 0.1 M Na_2_CO_3_, pH 9.0) for 30 min at 37°C. Then, yeast cells were washed in PBS and macrophages were infected at MOI 5 for 6 hours. Afterwards macrophages were washed with PBS, lysed with 0.5 % Triton™-X-100 for 15 min. Released yeast cells were washed with PBS, with 2 % BSA in PBS, and counterstained with 50 µg/mL Alexa Fluor 647-conjugated concanavalin A (ConA) (Molecular Probes) in PBS at 37 °C for 30 min. The ConA-AF647-stained yeast cells were washed with PBS and fixed with Histofix (Roth) for 15 min at 37°C. As FITC is not transferred to daughter cells, differentiation of mother and daughter cells was possible: The ratio of FITC positive and negative yeast cells was evaluated by flow cytometry (BD FACS Verse^®.^ BD Biosciences, Franklin Lakes (USA)) counting 10,000 events. Data analysis was performed using the FlowJO^™^ 10.2 software (FlowJO LLC, Ashland (USA)). The gating strategy was based on the detection of single and ConA positive cells and exclusion of cellular debris.

For quantification of intracellular replication by fluorescence microscopy cells were fixed with Histofix (Roth) after incubation with macrophages and stained 30 min at 37°C with 25 µg/ml ConA-AF647 (Molecular Probes) to visualize non-phagocytosed yeast cells. Then they were mounted cell side down in ProLong Gold antifade reagent (Molecular Probes). As FITC is not transferred to daughter cells, differentiation of mother and daughter cells was possible and intracellular replication was observed by fluorescence microscopy (Leica DM5500B and Leica DFC360).

### Competition assay

This experiment was adapted from Graf et al., 2019 (Graf et al., 2019). A431 vaginal epithelial cells (Deutsche Sammlung von Mikroorganismen und Zellkulturen DSMZ no. ACC 91) were routinely cultivated in RPMI1640 media with L-glutamine (Thermo Fisher Scientific) containing 10 % heat-inactivated fetal bovine serum (FBS; Bio&SELL), at 37°C and 5% CO2 for no longer than 15 passages. For detachment, cells were treated with Accutase (Gibco, Thermo Fisher Scientific). For use in experiments, the cell numbers were determined using a Neubauer chamber system and seeded in a 6-well-plate (4×10^5^ cells/well) for 3 days. For infection experiments, the medium was exchanged by fresh RPMI1640 without FBS. *L. rhamnosus* (ATCC 7469) was grown in MRS broth for 72 h at 37°C. Before infection, bacteria were harvested, washed with PBS and adjusted to an optical density OD_600_ of 0.2 (∼1×108 CFU/ml) in RPMI1640. Then, one third of the total volume of the well of a 6-well-plate was inoculated for 18 h prior infection with *C. glabrata*. These wells were colonized with *C. glabrata* wild type and mutant separated or mixed in equal cell number to a final MOI of 1 for 24 h. The same settings were established in the absence of bacteria as controls. Additionally, in some wells infected with both strains in the presence or absence of bacteria, a final concentration of 8 ng/ml of fluconazole was added. Fluconazole was dissolved in DMSO and it was ensured that the final percentage of the organic dissolvent in the wells was bellowed 0.1%.

After 24 h, supernatants and attached cells were collected and vortexed thoroughly. Vaginal cells were treated with 0.5 % Triton^™^-X-100 for 5 min to lyse them and release adherent fungal cells. Samples were diluted appropriately with PBS. The diluted samples were plated on YPD plates with 1× PenStrep (Gibco, Thermo Fisher Scientific) and incubated at 37°C for 1-2 days until adequate growth for determining the CFUs was reached.

### Sequencing

For the DNA extraction, the strains were grown in YPD cultures for 16 h at 37°C and 180 rpm, and the following protocol was implemented to isolate DNA of high quality. The cultures were centrifuged for 5 min at 4000 rpm. The pellet was suspended in sorbitol 1 M and centrifuged for 2 min at 13000 rpm. Then the pellet was resuspended in SCEM buffer (1M Sorbitol; 100mM Na-Citrate pH 5,8; 50mM EDTA pH 8; 2% ß-Mercaptoethanol and 500 units/ml Lyticase (MERCK)) and incubated at 37°C for 2 h. Afterwards the samples were centrifuged at 5 min at 13000 rpm and the pellet was resuspended in proteinase buffer (10mM Tris-CL pH 7,5; 50mM EDTA pH 7,5; 0,5% SDS and 1mg/ml Proteinase K), incubated at 60°C for 30 min. Phenol:Chloroform:Isoamylalkol 25:24:1 was added after the incubation and the samples were vortexed for 4 min. Then they were centrifuged for 4 min at 13000rpm. Then the aqueous phase was transfer to a new tube and 1:1 volume of cold isopropanol was added. Samples were centrifuged for 15 min at 13000 rpm. The pellets were washed with 70% ethanol once and centrifuged again for 3 min at 13000 rpm. After drying, the pellet was resuspended in water and RNase. The genomic DNA was stored in −20°C until sequencing. The sequencing of the clinical strains was done by the company GENEWIZ, using Illumina NovaSeq 2×150 bp sequencing and 10M raw paired-end reads per sample package. Additionally, paired-end reads for non-petite clinical isolates were obtained from a previous study (NCBI SRA project SRP099102 (Carrete et al., 2018)). All reads were aligned to the *C. glabrata* reference genome (version s03-mo1-r06, (Dujon et al., 2004)) using bowtie2 version 2.4.1. Variants were called from the resulting alignments using the callvariants script in bbmap (version 38.44) (SOURCEFORGE, 2014) with standard parameters. The resulting variant files were applied to the reference genome by the bcftools consensus function (version 1.10.2) (GitHub, 2019).

### In silico analysis and statistics

All the results were obtained from at least three biological replicates (indicated in figure legends). Mean and standard deviation of these replicates are shown. Experiments performed with MDMs were isolated from at least three different donors (see figure legends). Data were analyzed using GraphPad Prism 5 (GraphPad Software, San Diego, USA). The data were generally analyzed using a two-tailed, unpaired Student’s t test for intergroup comparisons, if not indicated otherwise.

## DATA AVAILABILITY

Raw sequencing data that support the findings of this study are available in the Sequence Read Archive (SRA) of the NCBI under the accession number PRJNA665484 (www.ncbi.nlm.nih.gov/sra/PRJNA665484).

## ACKNOWLEDGEMENTS

We thank Daniel Fischer, Marcel Sprenger, Franziska Pieper, Sophie Austermeier, and Stephanie Wisgott for their help and support during isolation of mBMDMs, and during isolation and cultivation of hMDMs. Further, we thank Franziska Pieper for her technical assistance in MIC_50_ analysis; Marina Pekmezovic and Marisa Valentine for their help with epithelial cell culture; Volha Skrahina for her technical assistance in creating the double mutant; Marina Pekmezovic for her help with microscopy imaging. We also thank Dominique Sanglard for providing clinical strains BPY40/41 and Grit Walther for providing clinical strains from the National Reference Center for Invasive Fungal Infections (NRZMyk). The auto-MACS system for magnetic isolation of human monocytes was provided by the research group Fungal Septomics.

